# RE2DC Resolves the Robustness–Efficiency Trade-Off in Single Particle Cryo-EM 2D Classification

**DOI:** 10.1101/2022.11.21.517443

**Authors:** Szu-Chi Chung, Hsin-Hung Lin, Tien-You Liu, Ting-Li Chen, Kuen-Phon Wu, Wei-Hau Chang, I-Ping Tu

**Author notes:** Correspondence (W.H.C.), (I.P.T.). Computational Precision Health, UCSF, San Francisco, CA 94143, U.S.A.

## Abstract

Single-particle cryo-electron microscopy (cryo-EM) increasingly generates millions of particle images, yet two-dimensional (2D) classification remains a major bottleneck because existing approaches balance computational efficiency against robustness to noise, outliers and structural heterogeneity. We introduce RE2DC (<u>R</u>obust and <u>E</u>fficient <u>2D</u> <u>C</u>lassifier), an algorithmic framework that resolves this trade-off through dynamic linear-time clustering, dimension-reduction multi-reference alignment, and offers real-time interactive *t*-SNE visualization. Rather than relying primarily on hardware acceleration, RE2DC reduces the computational cost of robust clustering and employs de-noised images for alignment, enabling efficient execution on standard multi-core CPUs. Across diverse benchmark datasets, RE2DC achieves class homogeneity comparable to ISAC while processing datasets three- to ten-fold faster per classification round than RELION. Notably, RE2DC resolves rare, structurally coherent particle populations, enabling detection of transient conformational intermediates and supporting near real-time cryo-EM analysis. By addressing algorithmic complexity, RE2DC establishes a general framework for robust, scalable analysis of massive and heterogeneous image datasets.

## Introduction

Cryogenic electron microscopy (cryo-EM) has transformed structural biology by enabling near-atomic to atomic resolution structure determination of bio-macromolecules in their native, unconstrained states [1,2]. This revolution has been driven by advances in direct electron detectors [3–6], modern image-processing pipelines [7–11] with robust algorithms [9-10,12-15] and automated microscope platforms capable of high-throughput, unceasing data acquisition [16–18]. Modern cryo-EM facilities routinely collect thousands of movie-micrographs daily, generating sub-millions of individual particle images. Unlike X-ray crystallography, single-particle cryo-EM preserves conformational heterogeneity, allowing structurally distinct functional states to be computationally separated rather than averaged away [19–21]. This unique capability has established cryo-EM as an indispensable tool for studying dynamic macromolecular machines, as demonstrated by the rapid structural characterization of the highly glycosylated SARS-CoV-2 spike protein and its emerging variants [22–25].

The unprecedented growth in data acquisition, however, has shifted the primary bottleneck from image collection to the extraction of discrete structural information from increasingly massive, noisy, and heterogeneous datasets [26]. A typical single-particle workflow incrementally recovers signal from images acquired under extremely low electron doses (typically 10–40 e^-^/Å^2^), resulting in signal-to-noise ratios often below 0.1 through successive stages of motion correction, contrast transfer function (CTF) estimation, particle picking, two-dimensional (2D) classification, and three-dimensional reconstruction [26,27] (Figure 1(a)). Among these steps, 2D classification occupies a pivotal position: it simultaneously aligns particle images, improves signal-to-noise ratios through class averaging, and groups images into distinct projection views. This step is the gatekeeper of data curation, determining which “good” particles proceed to 3D refinement and which “bad” particles (e.g., carbon edges, ice contamination, or damaged proteins) are discarded. Consequently, the quality of 2D classification directly dictates the accuracy and resolution of subsequent 3D reconstructions. Yet, reliable 2D classification remains a notorious user bottleneck, frequently requiring exhaustive parameter optimization, manual class curation, and multiple time-consuming rounds of computation.

**Figure 1.**
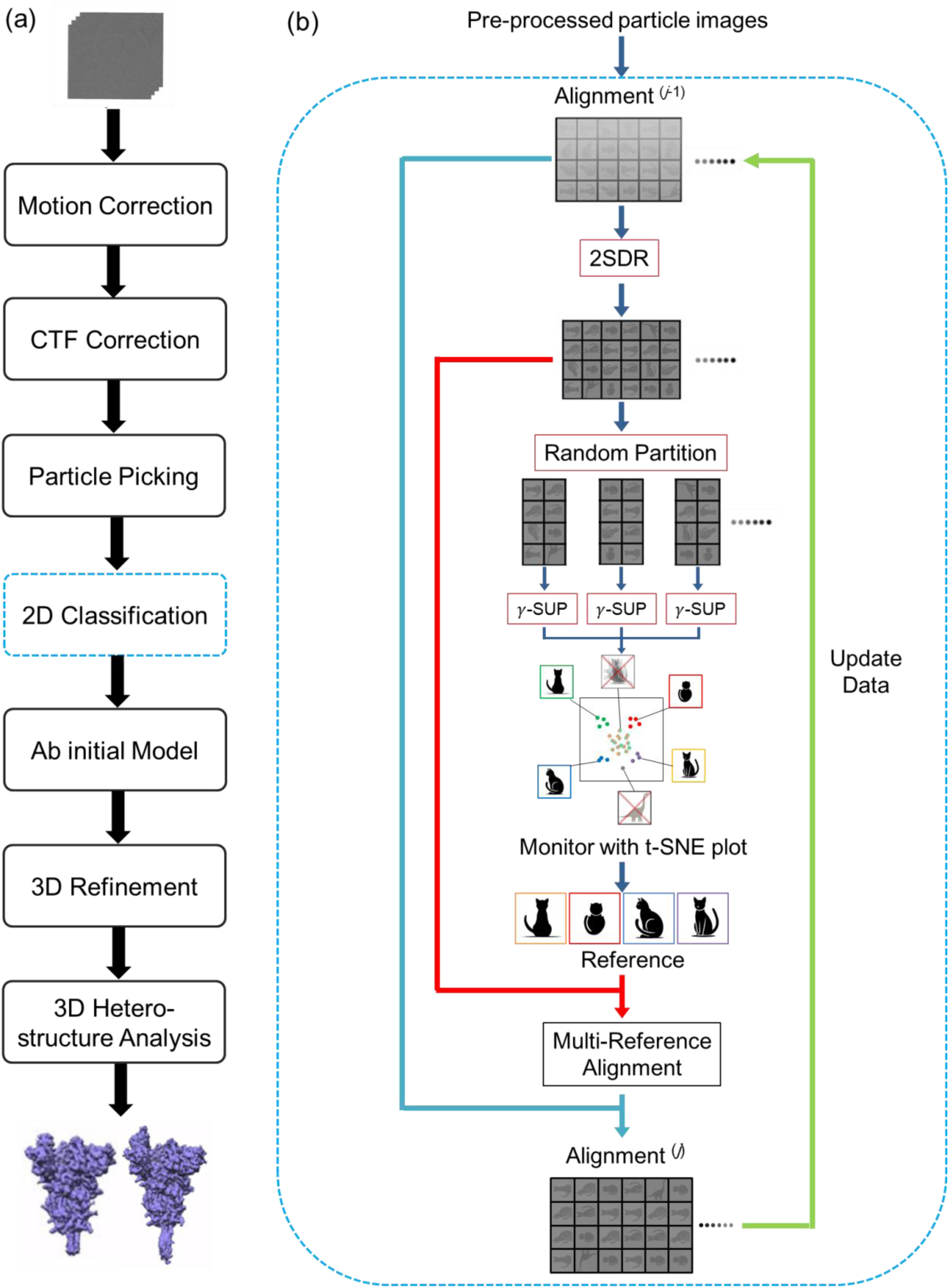
A standard workflow for processing single particle cryo-EM images. **(a)** Individual steps in the workflow from movie alignment (motion correction) to final density maps, as exemplified by a Covid-19 spike protein. **(b)** RE2DC algorithm for 2D classification. The iterative process is initialized with Pre-Pro preprocessing [35]. 2SDR, a two-stage tensor-structure dimension reduction method [38], is used to reduce images’ dimensionality and noise, allowing for their rapid and accurate alignment. For a large dataset, random partitioning is used to divide the data into several portions and processed in parallel via distributed computing; each partition then undergoes independent dynamic γ-SUP clustering. The resulting class average images are pooled to undergo another round of dynamic γ-SUP for cleaning up bad classes (omitted here but described in the algorithms in **Methods**). The resulting high-quality class average images are then used as references for multi-reference alignment. The estimated alignment parameters are used to update the pose parameters of the experimental images to generate better aligned images for the next iteration. In each iteration, t-SNE is utilized to monitor the distribution of images, each denoted by a colored dot.

This persistent bottleneck reflects a fundamental methodological challenge rather than a simple lack of raw computational power. Unlike conventional image clustering for simple and clear images, cryo-EM 2D classification is a non-trivial data science problem due to high level of noise and high image dimensionality [26]. Accurate classification requires the joint determination of image alignment pose parameters (translation and in-plane rotation) and particle class assignments [28,29], where the alignment was found to be highly error-prone under low SNR conditions [30]. Because alignment and clustering are strongly interdependent, errors in one process rapidly propagate to the other, creating a highly non-convex optimization landscape. Despite continuous algorithmic developments, existing 2D classification approaches are bound by a persistent trade-off—they can optimize for computational efficiency, or they can optimize for outlier-robustness and class homogeneity, but they have historically failed to achieve both. Widely adopted probabilistic approaches, such as the maximum-likelihood (ML) [14] and maximum a posteriori (MAP) frameworks implemented in RELION and CryoSPARC [9–10], prioritize computational efficiency enabled by GPU-acceleration [10,31] for routine data cleaning. By modeling particle images probabilistically, these methods use soft-assignment schemes that mitigate the risk of convergence to false extrema under extreme noise [32,33] to greatly advance cryo-EM structure determination. However, this soft assignment permits structurally distinct particles or low-confidence noise outliers to contribute to the same class average. This effect is often exacerbated in practice when users specify a modest number of classes (K) [34] to keep computational times manageable. Under these constraints, dominant structural views progressively absorb rarer or structurally distinct states—a phenomenon known as the “attractor effect” [13,34]—inflating within-class variance, reducing class homogeneity, and obscuring low-abundance sub-populations.

Conversely, validation-based approaches like ISAC (Iterative Stable Alignment and Clustering) [12] prioritize strict class homogeneity and outlier robustness. ISAC utilizes a constrained clustering framework that enforces equal-sized classes and aggressively rejects unstable or outlying particles across iterative validation loops. While this approach yields exceptionally homogeneous class averages capable of resolving subtle structure variation and conformational heterogeneity [35,36], its iterative validation scheme is incredibly expensive computationally. This limits its scalability, rendering it impractical for the massive million-particle datasets and real-time, on-the-fly feedback loops increasingly common in high-throughput facilities.

Here, we introduce **RE2DC** (**<u>R</u>**obust and **<u>E</u>**fficient **2D <u>C</u>**lassifier), a framework designed to break this efficiency–robustness trade-off by innovating both the underlying clustering mathematics and the dimensionality reduction scheme. RE2DC reformulates 2D classification as a robust clustering problem executed within an optimized, low-dimensional representation space (Figure 1(b)). The framework integrates three synergistic innovations (see **Methods**): **(i) Dynamic γ-SUP**—an automatic, scalable, linear-time (O(N)) descendant of the robust **γ**-SUP clustering algorithm [37] that models data as a mixture of q-Gaussian functions (q < 1) and utilizes robust **γ**-divergence to adaptively down-weights and rejects outlying noise particles without requiring the pre-specification of the class number (K), preserving highly imbalanced or rare structural populations; (ii) **Dimension-Reduction Multi-Reference Alignment (DRMRA)**—a rapid alignment engine that incorporates two-stage dimension reduction (2SDR) [38] to denoise images in a tensor-reduced space, drastically improving the accuracy and speed of rotational and translational alignment; (iii) **2SDR-accelerated t-SNE Visualization**—an enhanced implementation of t-SNE [39,40] in which the conventional PCA preprocessing is replaced by the computationally efficient and structure-preserving 2SDR dimensionality reduction [38]. This eliminates the computational bottleneck and mitigates potential data structure distortion associated with linear PCA projections [38,41], enabling high-fidelity embeddings to be continuously updated in sync with each RE2DC iteration rather than generated only after completion as commonly practiced by existing classification algorithms [42–44]. The resulting real-time visualization provides immediate feedback on the evolution of particle distributions and classification trajectories throughout the clustering process, unveiling the black box of a 2D classification process. In addition, we further introduced an interactive *t*-SNE (i-*t*-SNE) interface for RE2DC that transforms visualization from a passive display into an exploratory analysis tool, enabling users to identify ambiguous regions, examine class boundaries, and optionally perform user-guided refinement of particle groupings.

By avoiding the heavy, iterative multi-round optimization cycles of probabilistic frameworks and the expensive validation loops of hard-clustering methods, RE2DC converges to highly stable, homogeneous solutions with minimum number of iterations in a single classification round. We demonstrate the utility of RE2DC by benchmarking it against both ISAC and RELION using diverse, publicly available datasets. RE2DC matches or exceeds the extreme outlier-robustness of ISAC on a classic 70S ribosome dataset [9] while mirroring the actual experimental class size distribution. Scalability tests on massive, curated and uncurated datasets—including 80S ribosome (∼10^5^ particles) [EMPIAR-10028] [45] and TRPV channel datasets (∼0.9 times 10^5^ and 2.2 times 10^5^ particles) [EMPIAR-10005 & EMPIAR-10059] [46,47]—reveal that RE2DC achieves a three- to ten-fold speedup per-round compared to RELION, and processes over ∼3.8 times 10^5^ particles of a SARS-CoV-2 spike protein dataset [EMPIAR-10514] in 8.5 hours on a standard multi-core CPU and in less than 8 hours with GPU-acceleration [48], detecting subtle conformation movements in this protein including the domain for ACE2-receptor binding.

Crucially, we show that RE2DC’s unique combination of robustness and scalability enables the discovery of biologically vital, underrepresented structural states that are routinely lost to the attractor effect of conventional pipelines. Using RE2DC, we successfully identified an unexpected, highly transient oligomeric state in glutamine synthetase (GS) and resolved subtle structural intermediates in Rad51 filaments [49]. Fully integrated into ASCEP ecosystem (<u>A</u>ccelerated <u>S</u>tream <u>C</u>ryo-<u>E</u>M <u>P</u>rocessing) using Scipion architecture [50], RE2DC provides an accessible, high-performance standalone tool, or as an upstream pipeline component for supporting rapid structure determination workflows [51] including CryoSPARC and uncovering biological heterogeneity from the earliest stages of cryo-EM data processing.

## Results

### Dynamic γ-SUP Clustering is Fast, Accurate, and Attractor-Resistant

Because the computational complexity of the original γ-SUP algorithm scales quadratically as O(N^2^) (where N is the number of images) [37], applying it directly to large cryo-EM datasets is practically prohibitive. To overcome this computational bottleneck, we developed an advanced variant: dynamic k-nearest neighbor (kNN) γ-SUP, hereafter referred to as dynamic γ-SUP. By utilizing a kNN graph to approximate pairwise image similarities and dynamically adjusting the neighbor count k over successive iterations (**Methods**), dynamic γ-SUP successfully reduces the computational complexity to O(N).

To validate the performance of dynamic γ-SUP, we designed benchmark tests using two synthetic cryo-EM datasets, each containing 5,000 particle images with known ground-truth cluster labels and pre-aligned orientations: (i) Balanced Dataset: it comprises 50 equal-sized clusters (100 noisy images per cluster) with identical orientation and pose parameters; (ii) Unbalanced Dataset: it emulates experimental data heterogeneity with 10 large clusters (400 images each) and 40 small clusters (25 images each) (**Methods**).

To enable prompt visualization of the particle image distribution, we introduced 2SDR-preprocessed t-distributed Stochastic Neighbor Embedding (t-SNE) to project these high-dimensional particle images into a two-dimensional space (**Methods**). As shown in the t-SNE plots (Figure 2(a) & (b)), the balanced dataset is resolved into 50 equal-sized clusters, while the unbalanced dataset is resolved into 10 large and 40 small clusters, corresponding to the 50 distinct projection views. These results demonstrate that under non-crowded conditions, *t*-SNE projection faithfully presents 50 distinct projection views as 50 distinct clusters, with cluster sizes directly reflecting the number of contained particles. In addition to enabling prompt visualization of particle image distribution, 2SDR-preprocessed *t*-SNE seems to confer a superior data partition pattern than PCA-preprocessed *t*-SNE (Supplementary Figure 1).

**Figure 2.**
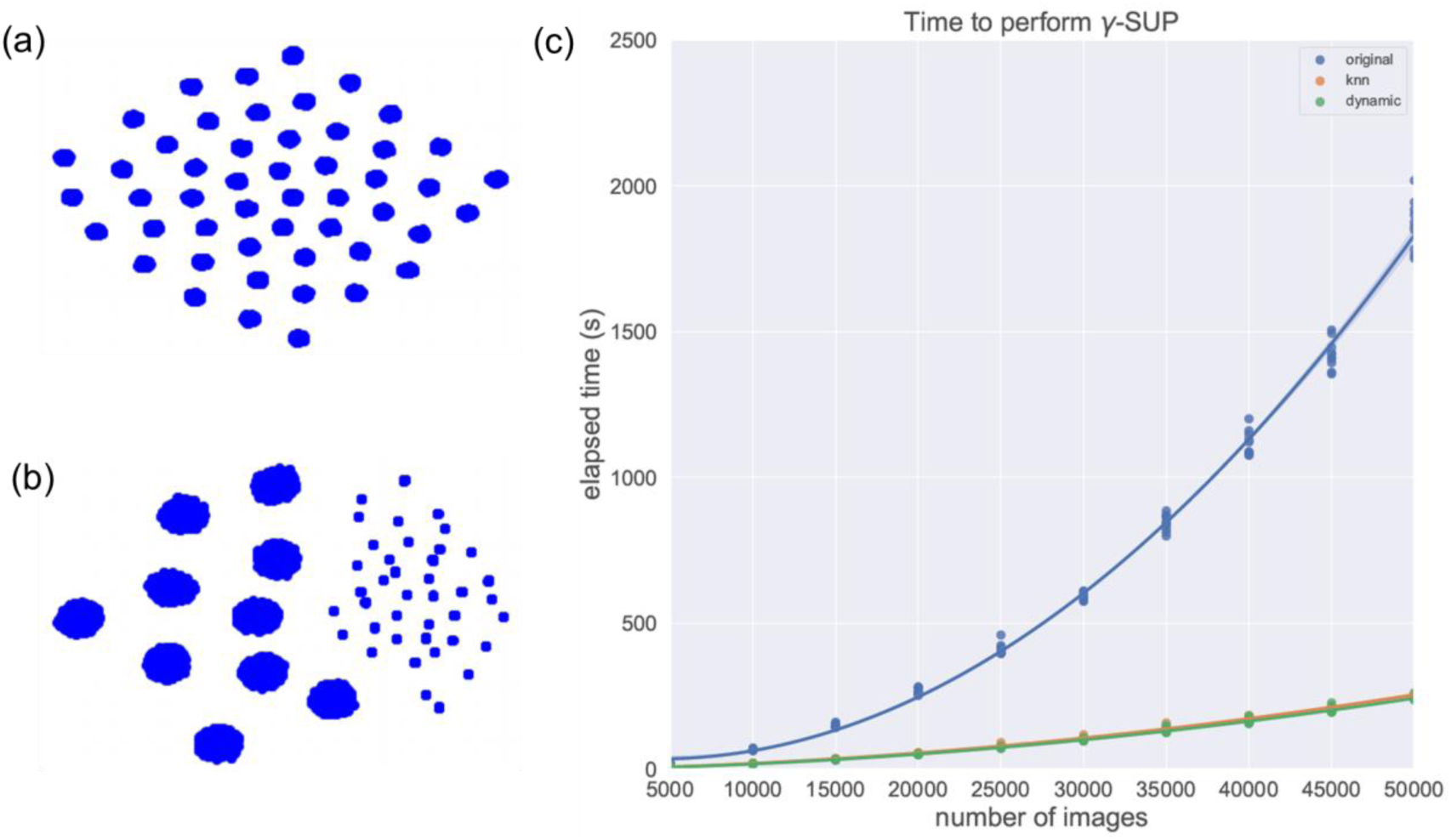
Tests for the performance of dynamic γ-SUP clustering using 70S ribosome synthetic data. **(a)** t-SNE visualization of clustering of a balance set of 50 view classes, each contains 100 particles. **(b)** t-SNE visualization of a unbalance set of 50 view classes, with each large class containing 400 particles while each small class containing 25 particles. **(c)** Comparison of the run time versus number of images for various γ -SUP clustering on an unbalanced set with SNR = 0.06 (original, kNN, dynamic γ–SUP; also see **Methods**).

We next evaluated how execution time scales with dataset size (N). As shown in Figure 2(c), the runtimes of both the kNN and dynamic γ-SUP algorithms scale linearly with N, consistent with their theoretical O(N) complexity, whereas the original γ-SUP algorithm scales quadratically (O(N^2^)). Remarkably, the dynamic γ-SUP completed clustering for a large dataset of 50,000 images in less than 250 seconds, demonstrating outstanding computational efficiency.

To evaluate clustering accuracy, we calculated the V-measure [52], which assesses predicted clusters against ground-truth labels (**Methods**). All three algorithms (original, kNN, and dynamic γ-SUP) achieved 100% accuracy (V=1.0) across a wide range of signal-to-noise ratios (SNRs) and hyperparameter settings for both the balanced and unbalanced datasets (Supplementary Figure 2). Notably, not a single particle from the small clusters was incorrectly absorbed into the larger clusters. This exceptional resistance to the “attractor effect” highlights the inherent outlier robustness of the γ-SUP architecture. Collectively, these findings demonstrate that dynamic γ-SUP is highly accurate, extremely fast, and scales seamlessly to massive datasets while retaining its robust outlier-rejection capabilities.

### RE2DC is a Robust and Efficient 2D Classifier Equipped with Fast, Interactive *t*-SNE

To adapt dynamic γ-SUP for experimental cryo-EM datasets where image alignment is required, we integrated it into a dimension-reduction multi-reference alignment (DRMRA) framework. This resulted in RE2DC (Robust and Efficient 2D Classifier; Figure 1(b)), which operates in an unsupervised, iterative manner. To determine the number of iterations required for RE2DC to reach convergence, we benchmarked it using a classic dataset of 5,000 handpicked 70S ribosome particle images provided by Haixiao Gao and Joachim Frank, monitoring both the 2D class averages and t-SNE spatial distributions at each iteration. This dataset is a standard historical benchmark, previously used to evaluate RELION 1.1 [9], the original γ-SUP [37], and the Pre-Pro preprocessing algorithm [35]. As shown in Figure 3(a), the RE2DC class averages stabilized within only three to four iterations, matching the point where the particle cloud in the corresponding t-SNE plot segregated into well-defined, isolated sub-clusters. This rapid convergence contrasts sharply with conventional methods that require dozens of iterations to converge as previously measured [35], underscoring the efficiency of our framework.

**Figure 3.**
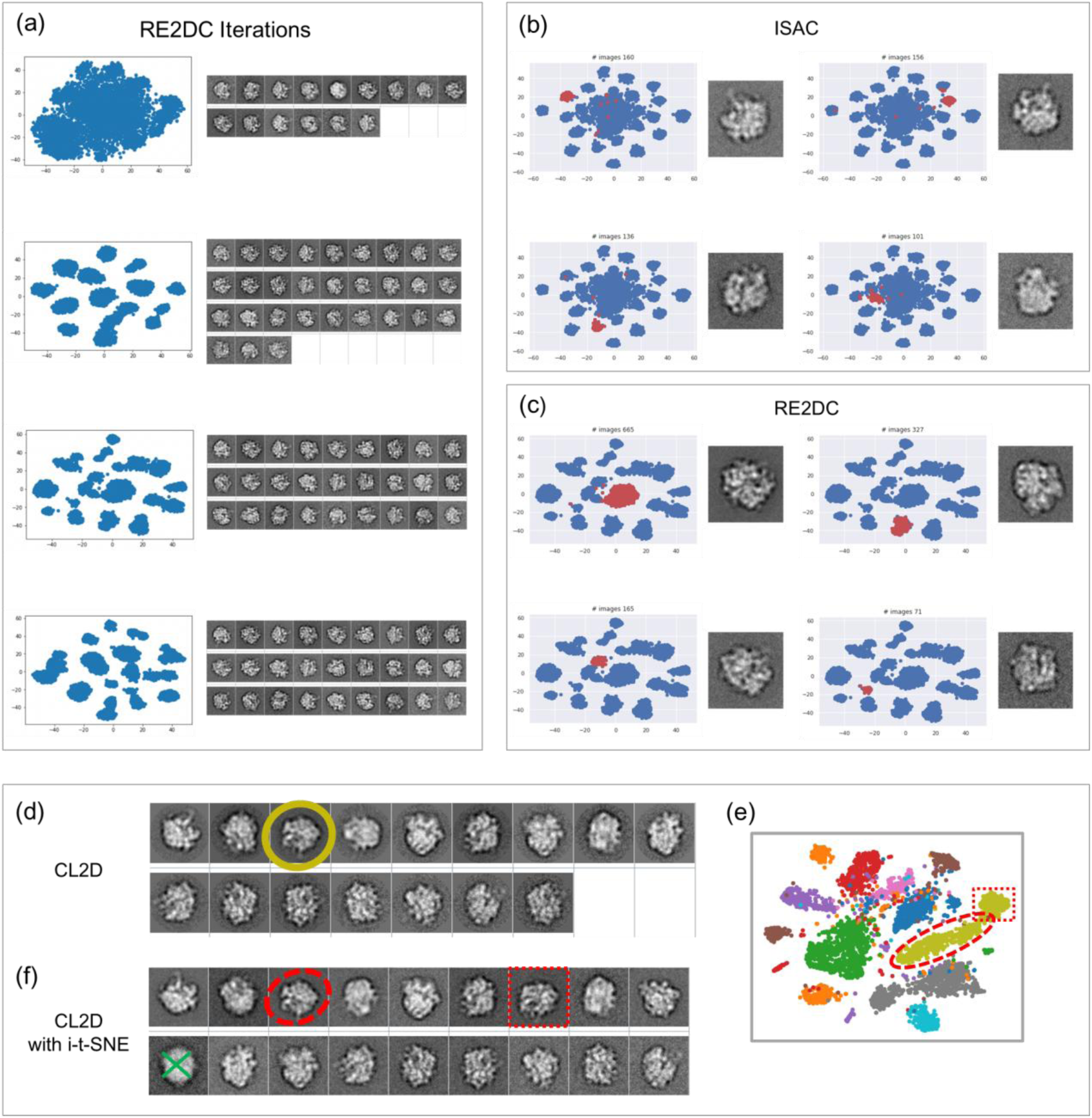
2D classification results of 70S ribosome with *t*-SNE visualization. **(a)** RE2DC 2D classification results visualized by *t*-SNE plots and class averages for the first four iterations. **(b)** ISAC results from the last iteration: four representative classes with *t*-SNE sub-plots and the class averages; the class members are denoted as red dots. **(c)** RE2DC results from the third iteration: four representative classes with t-SNE sub-plots and the class averages; the class members are denoted as red dots. **(d)(e)(f)** CL2D results: 16 class averages, the *t*-SNE plot, and the refined results with the aid of interactive *t*-SNE (i-*t*-SNE). In (e), an elongated cluster colored with earth color is observed, where it exhibits a sub-structure of a “sausage” connected to a small blob. The sausage and the small blob were lassoed respectively using i-*t-*SNE interface to generate two separate classes in (f).

In conventional workflows, t-SNE visualization relies on Principal Component Analysis (PCA) for initial dimension reduction. However, this PCA-preprocessing is faced with efficiency challenges for processing modern high-resolution cryo-EM particle images with large number of pixels (> 10^4^ pixels). To prevent this *t*-SNE visualization step from potentially hauling RE2DC’s iterations, we introduced a rapid t-SNE by combining a fast implementation [40] with Two-Stage Tensor-Structure Dimension Reduction (2SDR) [38] to replace the PCA dimension reduction in conventional t-SNE (**Methods**). Of note, conventional t-SNE or UMAP visualizations using PCA dimension reduction are prone to data structure distortions [38,41], and the associated high computational cost often restricts their use to final results rather than real-time monitoring during active classification iterations [42–44].

To demonstrate the outlier robustness of RE2DC, we compared it directly to ISAC, an established 2D classification method highly regarded for its ability to isolate impurities, reject outliers, and resolve conformational heterogeneity [12,36]. When previously applied to this 70S ribosome dataset, ISAC produced high-quality class averages representing diverse projection views [35]. Using a minimum class-size threshold of 200 particles, ISAC yielded 29 stable classes, retaining 3,875 particles (78% yield; Supplementary Figure 3).

The t-SNE projection of the particles aligned by ISAC in its final iteration revealed a distinct pattern (Figure 3(b), blue dots): approximately 20 small, isolated clusters distributed like satellites around a less-resolved central core. As expected due to ISAC’s equal-size K-means clustering constraints, these satellite classes each contained fewer than 200 particles [12]. However, projecting the actual members of each finalized ISAC class back onto the t-SNE space (Figure 3(b) & Supplementary Figure 3, red dots) revealed that while the majority of class members concentrated into single clusters, several particles were scattered far across the map. This indicates that even highly regarded robust classifiers like ISAC are not entirely free from class impurities.

Compared to ISAC, RE2DC similarly yielded 27 high-quality classes, but retained 4,667 particles (a 93% yield). The t-SNE pattern for RE2DC (Figure 3(c) and Supplementary Figure 4: *blue dots*) was fundamentally different. Rather than forcing equal-sized groups, the RE2DC classes showed a realistic range of sizes (from 71 to 665 particles, Figure 3(c) & Supplementary Figure 4: *blue dots*). This unequal distribution is a much more faithful representation of experimental cryo-EM data, where preferred particle orientations are common due to interactions at the air-water interface [53]. Furthermore, the t-SNE spatial distribution of aligned particles (blue dots) matched the class assignments made by dynamic γ-SUP (red dots) with extremely high fidelity. This demonstrates that RE2DC is highly effective at grouping physically similar, aligned particles while achieving superior outlier rejection and higher particle yields than ISAC.

Additionally, we observed that t-SNE projections can highlight discrepancies between spatial clusters of aligned images and the actual assignments made by a 2D classification algorithm (Supplementary Figure 5). This indicates that t-SNE can serve as a diagnostic and refinement tool. To demonstrate this possibility, we used CL2D [13] to classify the 70S ribosome dataset under sub-optimal conditions, targeting only 16 classes, far below the natural number of projection views. Intriguingly, the resulting *t*-SNE plot revealed that several CL2D classes contained hidden sub-structures; for example, the third class appeared as a “sausage-shaped” cluster connected to a smaller, distinct blob. To resolve them interactively, we developed interactive *t*-SNE (i-*t*-SNE) (**Methods**), allowing a user manually lasso and extract these sub-structures (e.g., separating the sausage cluster from the adjacent blob) for refining this sub-optimal CL2D class into two highly homogeneous subsets (Figure 3(f)). This interactive tool is particularly valuable for investigating crowded areas of t-SNE maps where automated classifiers struggle to draw clean boundaries (Supplementary Figure 6).

### RE2DC Uncovers a Hidden Quaternary State of Escherichia coli Glutamine Synthetase

Glutamine synthetase (GS) is a ubiquitous metabolic enzyme that catalyzes the ATP-dependent condensation of glutamate and ammonia to form glutamine. While eukaryotic GS typically assembles into octamers or decamers, prokaryotic GS variants predominantly form dodecamers or hexamers [54]. Although high-resolution structures of GS from various organisms have been solved via X-ray crystallography, Escherichia coli GS has historically resisted crystallization. Early negative-stain electron microscopy of E. coli GS revealed co-existing dodecamers alongside “broken” oligomeric forms [55]. We hypothesized that the physical fragility of E. coli GS leads to severe compositional heterogeneity in solution, which has historically hindered crystal growth.

To test this hypothesis, we subjected purified E. coli GS to single-particle cryo-EM. We collected a novel dataset on a Titan Krios microscope and picked 94,154 particles, which we then classified using RELION, ISAC, and RE2DC. As shown in Figure 4(a)-(c), all three methods generated similar 2D class averages for the dominant populations: clear “end-on” views (6-fold symmetric rings) and “side” views showing the characteristic double-layered disc of a dodecamer (12-mer). The massive over-representation of end-on views highlights a severe preferred-orientation issue [53], which typically makes high-resolution 3D reconstruction extremely challenging [56].

**Figure 4.**
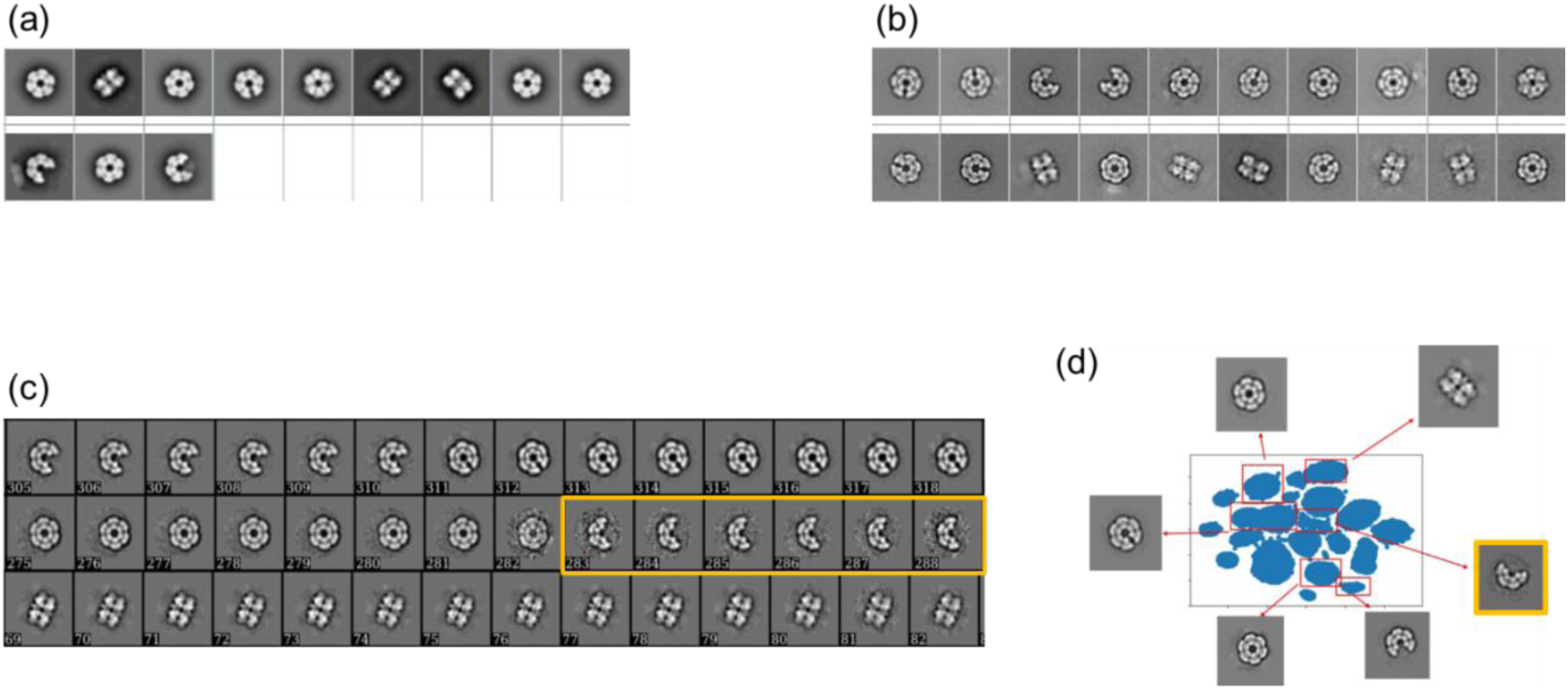
2D classification results of *E. coli* glutamine synthetase (GS) particle images from different methods. **(a)** Results from RELION. **(b)** Results from ISAC. **(c)** Results from RE2DC (see **Methods**); the yellow boxes highlight a peculiar broken GS form (8-mer). **(d)** *t*-SNE plot for the particles aligned by RE2DC with a few representative clusters investigated by i-*t*-SNE. The orange box highlights the peculiar broken form (10-mer).

A closer inspection of these end-on views confirmed that the *E. coli* GS sample is highly heterogeneous. The major populations across all methods included intact and notched dodecamers and a broken form lacking two subunits (a 10-mer). Strikingly, however, RE2DC uniquely resolved a crescent-shaped form lacking four subunits (an 8-mer) (Figure 4(c) & Supplementary Figure 7, highlighted in yellow). This rare, highly fragile state was nearly completely missed or merged into broader classes by RELION, ISAC, and CryoSPARC, in particular when pre-specified class parameters optimized for computational efficiency were used (Supplementary Figure 8-10).

To verify that this crescent-shaped state was physically real and not an algorithmic artifact, we generated a *t*-SNE plot for the RE2DC aligned particles. By averaging the data points within the resolved *t*-SNE clusters using i-t-SNE, we confirmed that these crescent-shaped particles (8-mer) occupy a distinct, smaller cluster near the center of the distribution (Figure 4(d)). These results also confirmed the physical existence of the 12-mer, 10-mer, in addition to the newly uncovered 8-mer crescent state, demonstrating the power of RE2DC in resolving rare and highly fragile macromolecular states.

### RE2DC is Highly Efficient for Curating Massive, Heterogeneous Datasets

To evaluate the scalability and speed of RE2DC on large-scale experimental data, we benchmarked it against three datasets of increasing size and complexity: (I) 80S Ribosome [EMPIAR-10028]: 105,247 pre-curated particle images of Plasmodium falciparum 80S ribosome bound to the drug emetine [45]. Because the drug restricts the ratcheting motion between the 40S and 60S subunits, the dataset’s heterogeneity is primarily spatial (projection views) rather than conformational; (II) **NC-TRPV1 [**EMPIAR-10005]: 88,915 non-curated, highly heterogeneous TRPV1 ion channel particles [46]; (III) **NanoD-TRPV1 [**EMPIAR-10059]: 218,805 non-curated TRPV1 particles embedded in lipid nanodiscs [47]. The TRPV1 datasets are highly challenging because the channel is relatively small, highly dynamic, and surrounded by dense, disordered detergent micelles or lipid nanodiscs. The original authors had to use extensive, multi-round 2D and 3D classification to pull out homogeneous subsets, ultimately achieving resolutions of 3.4 Å (NC-TRPV1) and 2.9 Å (NanoD-TRPV1).

For a head-to-head comparison on the 80S ribosome dataset, we configured RELION, ISAC, and RE2DC to target 500 classes in a single round. Because this is a highly curated dataset, we expected particle yields to approach 100%. To run RE2DC without overcrowding the *t*-SNE projections, we split the dataset into 20 subsets and processed them in parallel using a distributed computing strategy (Figure 1(b); **Methods**). While all three methods achieved nearly 100% particle yields, they differed dramatically in class quality and runtime: **RELION** generated only 266 total classes, of which only 141 were of high quality (Supplementary Figure 11 & 12). Its execution took 62 hours on CPU and 30 hours with GPU acceleration. **ISAC** successfully resolved the data into 514 high-quality classes (Supplementary Figure 11 & 13), taking 124 hours on CPU and 14 hours using its newly released GPU-accelerated version [57]. **RE2DC** resolved the data into 542 clean classes (Supplementary Figure 11 & 14). On CPU using our distributed computing framework, RE2DC completed the entire run in just 2.5 hours—a 10-fold speedup over GPU-accelerated RELION and a 50-fold speedup over CPU-based ISAC. Incorporating GPU acceleration further reduced RE2DC’s runtime to just 2.0 hours (Table 1 & Supplementary Table 1).

**Table 1.**
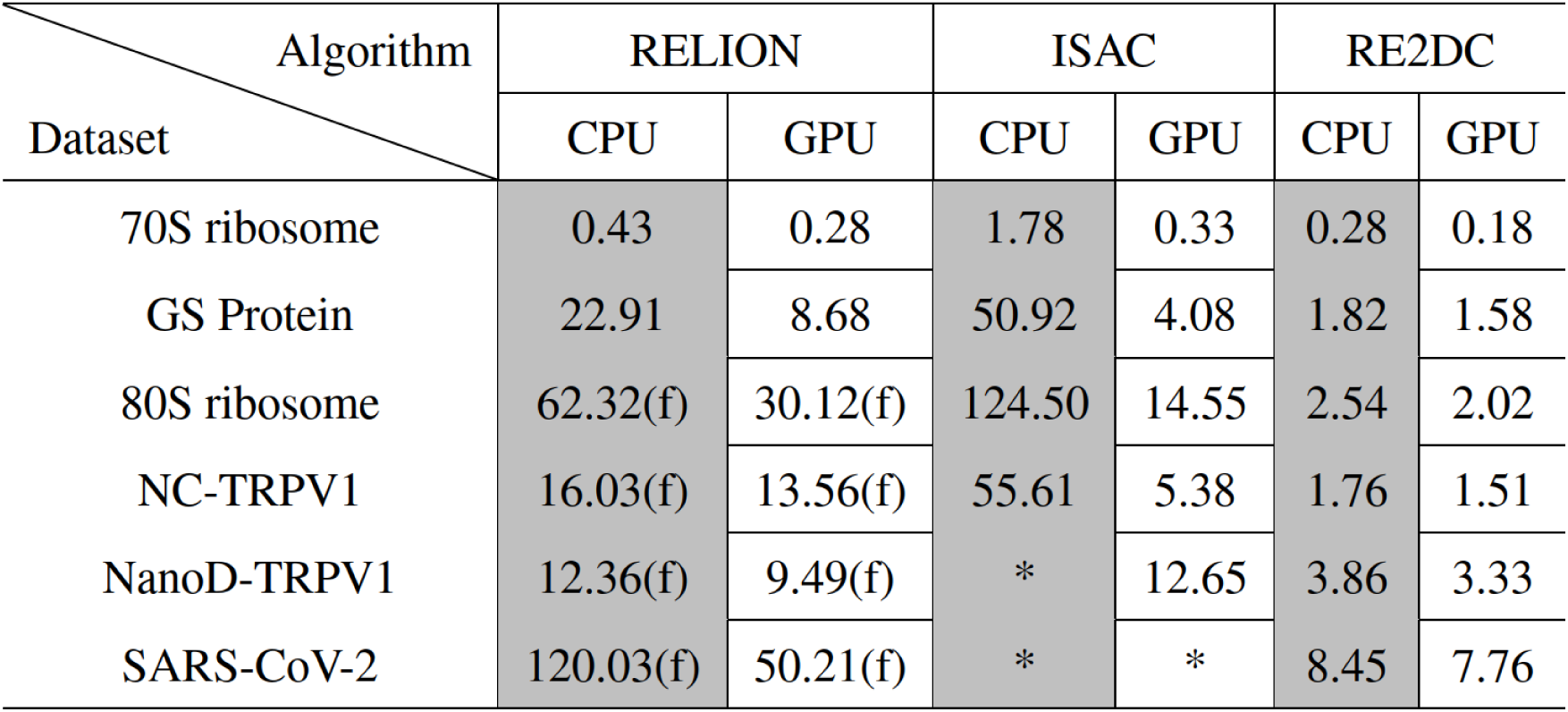
Execution time analysis for all datasets. To perform RE2DC, partition sizes are set as follows: 5,000 for the 70S ribosome, 80S ribosome, and NC-TRPV1, 10,000 for GS and NanoD-TRPV1, and 20,000 for SARS-CoV-2. Dynamic γ-SUP is utilized during the clustering step in RE2DC. We use 44 Message Passing Interface (MPI) tasks and a single GPU. Execution time represents the average from 5 measurements for the 70S ribosome and 3 measurements for all others. Note that the time for data format conversion on the platform is excluded from the measurement. Please see Supplementary Table 1 for more details about other settings. “(f)” denotes “fast-subset” option on RELION. An asterisk (*) indicates that the run had failed.

| Dataset \ Algorithm | RELION |  | ISAC |  | RE2DC |  |
| --- | --- | --- | --- | --- | --- | --- |
|  | CPU | GPU | CPU | GPU | CPU | GPU |
| 70S ribosome | 0.43 | 0.28 | 1.78 | 0.33 | 0.28 | 0.18 |
| GS Protein | 22.91 | 8.68 | 50.92 | 4.08 | 1.82 | 1.58 |
| 80S ribosome | 62.32(f) | 30.12(f) | 124.50 | 14.55 | 2.54 | 2.02 |
| NC-TRPV1 | 16.03(f) | 13.56(f) | 55.61 | 5.38 | 1.76 | 1.51 |
| NanoD-TRPV1 | 12.36(f) | 9.49(f) | * | 12.65 | 3.86 | 3.33 |
| SARS-CoV-2 | 120.03(f) | 50.21(f) | * | * | 8.45 | 7.76 |

Next, we evaluated the two non-curated, highly heterogeneous membrane protein datasets. For the NC-TRPV1 dataset, RELION took over 13 hours to complete classification, recovering roughly 60% of the particles. RE2DC completed the task in under 2 hours (1.7 hours on CPU, 1.5 hours on GPU) while achieving a superior particle yield of approximately 70% (Supplementary Figure 15-16). For the massive NanoD-TRPV1 dataset, RELION required nearly 10 hours and yielded 70% of the particles. In contrast, RE2DC completed the classification in under 4 hours (3.8 hours on CPU, 3.3 hours on GPU), recovering 73% of the particles (Supplementary Figure 15 & 19). This yield is roughly double the number of clean particles isolated by the original authors through their laborious multi-round 2D/3D sorting pipeline, yet it yielded a 3D reconstruction of comparable resolution (Supplementary Table 1). These results demonstrate that RE2DC is faster and more effective than conventional tools for cleaning up large, noisy, and highly heterogeneous membrane protein datasets in a single round.

### RE2DC Accelerates SARS-CoV-2 Spike Data Processing and Flags Highly Flexible Structural Domains

The SARS-CoV-2 spike protein (2019-nCoV S) is a trimeric fusion machine. The first single-particle cryo-EM dataset of this spike stabilized in its pre-fusion state [22] was famously difficult to process due to severe structural heterogeneity. The original authors had to perform extensive, multi-round 2D and 3D classification to narrow down their initial pick of approximately 600,000 particles to 262,732 high-quality particles [22]. 3D classification of these particles showed that the 21 kDa receptor-binding domains (RBDs) are highly dynamic, adopting a mixture of “1-RBD-up/2-RBD-down” conformations (Figure 5(a)) and causing severe local density blurring. Recently, Carazo’s group re-analyzed these raw movie frames [EMPIAR-10514] [58] using advanced continuous flexibility analysis, further resolving the “up” conformations into distinct open and closed intermediate states.

**Figure 5.**
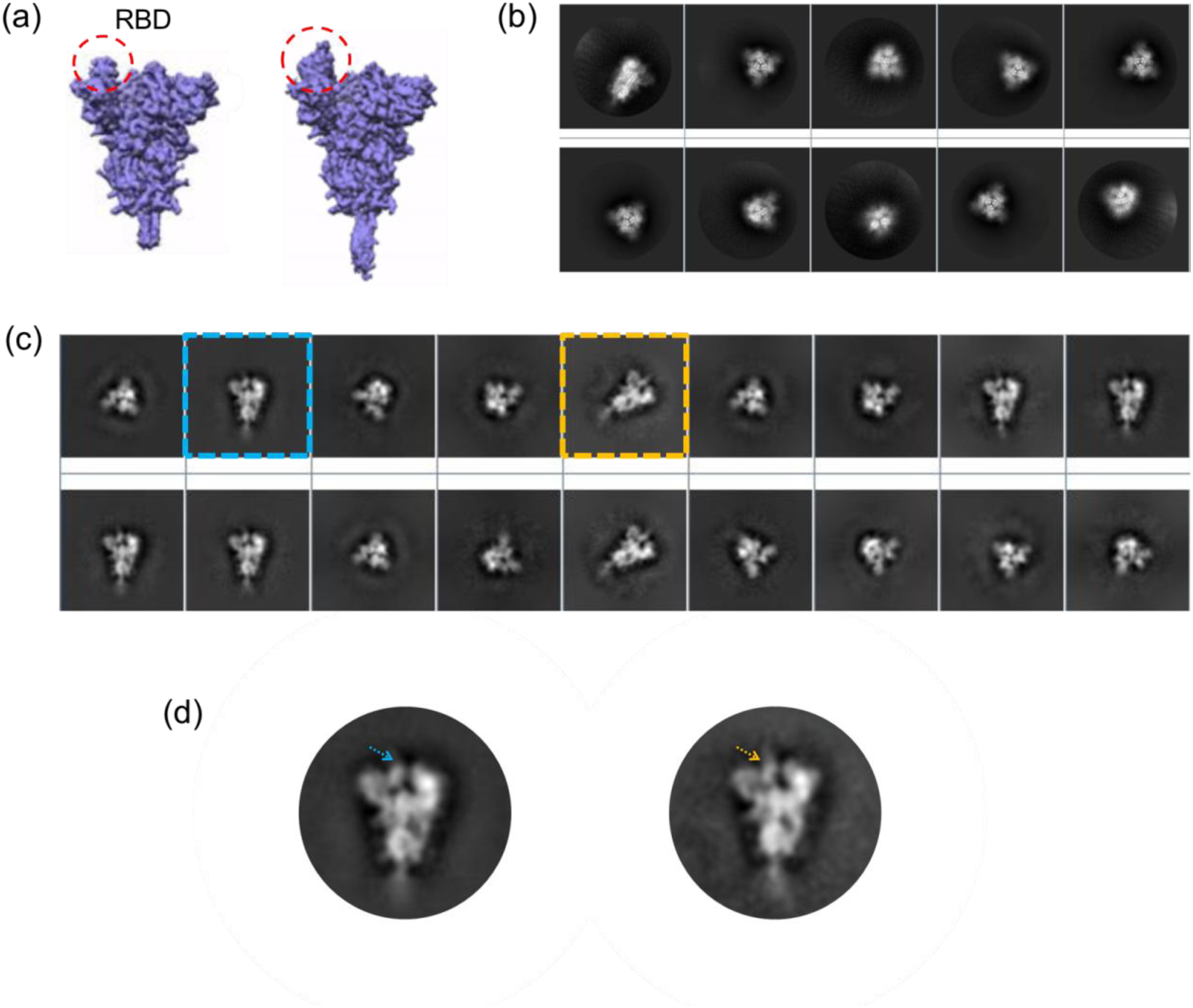
2019-nCoV Spike particle 2D classification analysis. **(a)** Two structures of the S protein obtained from the dataset [EMPIAR-10514] using CryoSPARC’s 3DVA analysis, as showcased on CryoSPARC website. The ACE2-recpetor binding domains (RBD) are highlighted. **(b)** 2D classes from RELION. **(c)** 2D classes from RE2DC. Two side-view classes that differ in the angle of S protein axis are boxed in blue and orange respectively. **(d)** The side-view corresponding to that boxed in blue in **(c)** is rotated so that the S protein axes for the two classes are in the same direction. The blue (orange) arrow points to the RBD domain.

To test RE2DC’s speed and real-world utility on a high-priority viral target, we downloaded the 3,511 raw cryo-EM movies [EMPIAR-10514]. Following the preprocessing protocols described by Carazo and colleagues [58], we extracted 375,067 particles—the largest and most heterogeneous dataset in this study—and compared the performance of RELION, ISAC, and RE2DC: (I) RELION completed 2D classification in 50 hours, recovering 235,300 good particles (63% yield); (II) ISAC failed to handle the massive dataset and terminated prematurely due to memory/scaling limitations; (III) RE2DC successfully completed the classification in just under 9 hours (8.5 hours on CPU, 7.8 hours on GPU), recovering 221,916 high-quality particles (59% yield).

Using these RE2DC-selected particles, we obtained a 3.44 Å 3D density map in CryoSPARC (Supplementary Table 1), well matching the resolution published by the original authors [22]. Given that raw data collection for these 3,511 movies takes roughly 10 hours on modern high-throughput instruments [16–18], and subsequent 3D analysis in CryoSPARC takes a few hours [10], replacing conventional 2D classification with RE2DC makes it entirely feasible to go from raw sample on the microscope to a completed 3D structure in less than 24 hours.

Beyond its speed, RE2DC revealed important structural insights directly from 2D classification. While the RELION class averages were heavily dominated by top-down end-on views with only a single side-view class (Figure 5(b) & Supplementary Figure 20), the RE2DC class averages showed a much more balanced distribution of top and side views (Figure 5(c) and Supplementary Figure 21).

Intriguingly, RE2DC separated the side-view particles into two highly similar yet distinct classes rather than merging them. One class featured an S-protein axis pointing vertically upward (Figure 5(c), blue box), while the other showed the axis tilted at a ∼45-degree angle (Figure 5(c), orange box). A close comparison of these two side-view classes (Figure 5(d)) rotated into a nearly identical coordinate system revealed subtle but distinct variations in density. In particular, we observed different patterns of density blurring around the RBD region (Figure 5(d), blue and orange arrows). This 2D observation perfectly mirrors the continuous RBD flexibility later resolved by complex 3D continuous flexibility analysis [58]. This demonstrates that RE2DC’s high-fidelity 2D classification can flag highly flexible, biologically important regions of a macromolecule early in the processing pipeline, well before initiating computationally expensive 3D heterogeneity analysis.

## Discussion

Single-particle cryo-EM data processing has long been constrained by a persistent methodological trade-off: pipelines must sacrifice computational speed for outlier robustness, or sacrifice sensitivity to rare structural states for scalability. Conventional maximum-likelihood (ML) and maximum a posteriori (MAP) algorithms rely on soft assignments that accelerate convergence but inadvertently induce the “attractor effect,” where low-abundance or fragile conformations are absorbed into dominant projection views. Conversely, rigorous hard-clustering and validation-based frameworks preserve extreme homogeneity at the cost of prohibitive computational overhead. RE2DC breaks this trilemma of robustness-efficiency-scalability by introducing linear-time dynamic γ-SUP clustering and combining it with a two-stage tensor-structure dimension reduction (2SDR) alignment engine. More importantly, RE2DC achieves this efficiency by reducing computational complexity rather than relying on hardware acceleration. Its performance arises from two complementary algorithmic strategies—linear-time robust clustering and aggressive data-dimensionality reduction—that are broadly applicable to large-scale data analysis beyond cryo-EM. Consequently, RE2DC achieves competitive performance on conventional CPU hardware, while GPU acceleration provides only incremental improvements because the dominant computational bottlenecks have already been removed algorithmically.

### Resolving the Robustness–Scalability Trade-Off to Preserve Structurally Coherent Rare States during 2D Classification

By combining robust q-Gaussian modeling (q < 1) with γ-divergence, RE2DC suppresses the influence of extreme noise/outliers while preserving structurally coherent particle populations, without requiring prior specification of the number of classes (K). This data-driven, attractor-resistant architecture allows small, structurally coherent particle sub-populations to persist alongside dominant orientations. As demonstrated across diverse experimental targets—from the recovery of a novel, highly transient crescent-shaped octameric intermediate in E. coli glutamine synthetase, and structural intermediates in human Rad51 filaments [49], to flagging continuous RBD domain flexibility in the SARS-CoV-2 spike protein—RE2DC enables underrepresented conformational states to remain distinguishable during 2D classification. Early preservation of these rare conformational states not only provides immediate mechanistic clues but also guides subsequent 3D reconstruction, reducing the need for extensive trial-and-error 3D classification while decreasing the risk that biologically meaningful particle populations are discarded during early image processing.

### High-Throughput Scalability and Potential Real-Time “Live2D” On-the-Fly Application

Beyond its robustness, RE2DC addresses the acute computational bottleneck created by modern high-throughput microscope platform generating tens of thousands of movies daily. By reducing algorithmic complexity from quadratic O(N^2^) [37] to linear O(N), RE2DC achieves a 3- to 10-fold speedup per round over RELION on massive, uncurated datasets while matching or exceeding particle yields. Notably, using RE2DC for particle curation normally requires only a single round.

Crucially, this combination of rapidity and stability may make RE2DC ideally suited for on-the-fly “Live2D” processing synchronized with live data acquisition. Platforms such as Warp [59], RELION [60], SIMPLE [61], and CryoSPARC Live perform real-time motion correction, CTF estimation, and particle picking, but on-the-fly 2D classification has traditionally lagged behind due to that the large particle numbers required for stable likelihood calculations are to be accumulated via a long microscope session. In contrast, RE2DC can process streaming batches (e.g., ∼5,000 particles) in about 10 minutes with GPU acceleration and under 20 minutes on a standard multi-core CPU. This capability, when implemented on a microscope workstation, would enable experimentalists to carry out on-the-fly 2D classification to continuously monitor data quality, evaluate orientation distributions, and diagnose severe preferred-orientation biases [62,63] early enough to allow a user to adjust acquisition strategies (e.g., applying sample tilt [64]) while the sample is still on the microscope.

### Dynamic, Interactive Diagnostics and Ecosystem Integration

Most conventional 2D workflows present classification as a static black box, yielding only discrete class averages. RE2DC transforms visualization into an active diagnostic tool by introducing 2SDR-accelerated *t*-SNE and embedding it directly into iterative classification loops. Because 2SDR bypasses linear PCA projections, it removes the associated computational overhead, allowing dynamic *t*-SNE updates to track alignment trajectories in real time. Furthermore, the interactive *t*-SNE (i-*t*-SNE) interface allows users to visually inspect crowded embedding regions, evaluate class boundaries, and manually extract hidden sub-populations from sub-optimal classifications.

RE2DC is designed to complement, rather than replace, existing 2D and 3D pipelines. Operating efficiently on accessible standard CPU hardware as well as GPU setups, RE2DC offers an energy-aware, scalable solution for massive data curation. Fully integrated into the Scipion/ASCEP ecosystems, RE2DC bridges raw data acquisition with advanced downstream continuous heterogeneity tools (such as HetSiren [65]), establishing a streamlined and robust foundation for high-resolution single-particle cryo-EM. It is our mission to ensure RE2DC is being continuously renewed, for which a new version has been edited in response to recent Scipion update and ready to replace the last version [66].

## Methods

Here, we summarize the conceptual framework for our methodological innovations. Detailed algorithmic implementations and data processing procedures are available in the **SI Methods**.

### Dynamic γ-SUP: a robust and scalable linear-time clustering method

Previously, γ-SUP was introduced as an accurate, noise- and outlier-robust clustering algorithm for analyzing cryo-EM particle images [37]. Unlike conventional model- or distance-based clustering methods, γ-SUP adopts a hybrid formulation that models data as a mixture of q-Gaussian functions (q < 1) and employs γ-divergence to measure similarities between data and model distributions. This enables γ-SUP to reject data points outside the effective support of a cluster while progressively down-weighting observations that deviate substantially from cluster centers. Furthermore, γ-SUP requires neither pre-specification of the number of clusters (K) nor random initialization, as every data point starts as a singleton cluster. This design avoids the random initialization pitfalls inherent to K-seed strategies, such as those used in RELION [9,34] or K-means-based approaches like ISAC [12]. While γ-SUP excels when analyzing datasets with numerous distinct projection views, heavy noise or outliers, or highly unbalanced cluster sizes [37], its application to massive datasets has been bottlenecked by three factors: (i) Computational complexity that scales quadratically with image count (O(N^2^)); (ii) The need for manual hyperparameter tuning by inspecting a plot to identify the transition point (see Fig. 5 in [37] & Supplementary Figure 22); (iii) The lack of built-in image alignment, requiring a separate alignment step. To address these limitations, we developed dynamic γ-SUP, a computationally efficient and scalable descendant of γ-SUP. Leveraging the inherent sparsity of the γ-SUP weight matrix (arising from the use of γ-divergence), we introduced a sparse approximation using a k-nearest neighbor (kNN) graph constructed via space-partitioning trees and hierarchical graphs [67,68]. This reduced the computational complexity of kNN γ-SUP from O(N^2^) down to O(N log N) or even O(N). We then made k dynamically adjustable across iterations to establish dynamic kNN γ-SUP, which we abbreviated as dynamic γ-SUP. To eliminate manual parameter tuning, we devised an automated, two-stage hyperparameter selection strategy: (i) Binary Search: Identifying a “non-trivial” region—avoiding states where all points collapse into a single cluster or remain as singletons; (ii) Grid Search: Scanning this region to maximize an objective target function (e.g., particle yield, number of reasonable-sized clusters, or a combination thereof).

### Dimension-Reduction Multi-Reference Alignment (DRMRA) framework

To incorporate image alignment into dynamic γ-SUP for real cryo-EM datasets, we developed the Dimension-Reduction Multi-Reference Alignment (DRMRA) framework and integrated it with dynamic γ-SUP (Figure 1(b)). DRMRA leverages 2SDR [38], a tensor-based method that efficiently achieves simultaneous image dimensionality reduction and denoising. Crucially, 2SDR denoising mitigates the risk of converging to false local minima during image alignment under high noise conditions—a known challenge in early 2D classification methods that let image noise less fettered [27–29]. Combining dynamic γ-SUP with DRMRA yielded RE2DC (Figure 1(b)), an unsupervised, scalable, and robust 2D classifier tailored for large-scale, heterogeneous cryo-EM datasets.

### 2SDR-accelerated *t*-SNE visualization

RE2DC includes a 2SDR-accelerated t-distributed stochastic neighbor embedding (*t*-SNE) module to offer real-time data visualization for tracking each iteration of a 2D classification process. Conventional *t*-SNE visualization cannot be used for 2D classification in real-time because it relies on PCA for pre-embedding dimension reduction, which incurs substantial computational overhead for processing high-dimensional cryo-EM images (>10^4^ pixels for a 100 times 100 box). By using 2SDR [38] to replace PCA for pre-embedding dimension reduction alongside FFT-accelerated interpolation [40], our implementation allows *t*-SNE visualization to keep pace with each RE2DC iteration. Additionally, we developed an interactive interface (i-*t*-SNE) to enable real-time diagnosis and interactive refinement of 2D classification results.

### ASCEP

RE2DC has been integrated into ASCEP (<u>A</u>ccelerated <u>S</u>tream <u>C</u>ryo-<u>E</u>M <u>P</u>rocessing) using the architecture of Scipion [51] and will be made available as part of ASCEP plugin of Scipion V3.0 (http://scipion.i2pc.es). This allows RE2DC to be connected virtually to most Cryo-EM image processing pipelines, including CryoSPARC [10], for supporting downstream 3D structure analysis.

## Data availability

The 70S ribosome can be found in EMDB test image data. 80S ribosome is from EMPIAR-10028. TRPV1 channel is from EMPIAR-10005. TRPV1 embedded in nanodisc is from EMDB EMPIAR-10059. SARS-CoV-2 stabilized spike in the prefusion state is from EMDB EMPIAR-10514.

## Code availability

The code will be deposited in GitHub as soon as this manuscript is accepted by a peer-reviewed journal.

## Competing interests

The authors declare no competing interests.

### Acknowledgement

This work was supported by Academia Sinica Grants: [AS-GCS-108-08] and [AS-IA-110-M05] to I-P.T.; Academia Sinica SUMMIT Projects [AS-SUMMIT-107], [AS-SUMMIT-108], [AS-SUMMIT-109] to W.-H.C.; and by the Ministry of Science and Technology Taiwan: [MOST 106-2118-M-001-001-MY2] to I-P.T.; [MOST 106-2321-B-001-050-MY3] to K.-P.W.; and [MOST 110-2118-M-110-003-MY2] to S.-C.C.. The authors are thankful to Academia Sinica Cryo-EM Center [AS-CFII-108-110] for supporting cryo-EM data collection. The authors are indebted to I-P.T.’s group members: Hui-Lun Shiao and Yu-Hsiang Lien for updating ASCEP, and Szu-Han Lin for proofreading the manuscript with special comments on the figures.

## Author Contribution

I-P.T. and W.-H.C. conceived the conceptual framework. S.-C.C. designed the computation architecture and performed the coding with conducting the classification experiments. T.-L.C. contributed to the conceptual development of the dynamic γ-SUP algorithm. K.-P.W. conducted GS protein purification, with its cryo-EM sample making, imaging and initial analysis. S.-C.C., H.-H.L., and W.-H.C. analyzed the algorithm performance and the classification results. S.-C.C., I.-P.T. wrote the early versions of the manuscript, with S.-C.C. and W.-H.C. editing the final version.

## Supplementary Information

## SI Methods

Existing 2D classification methods for cryo-EM images trades robustness for efficiency or vice versa, rarely achieving both. We here aim to build a 2D classification framework that is simultaneously robust and efficient. To do so, we chose a classical 2D classification framework that interweaves the image alignment and image clustering steps. A robust and efficient 2D classification method thus demands its sub-algorithms for alignment and clustering to bear these characters. With this in mind, we adopted γ-SUP clustering as the core algorithm for its proven robustness [1], and then developed complementary strategies to improve its efficiency. The robustness of γ-SUP is rooted in (i) use of the q-Gaussian distribution to define a rejection boundary and γ-divergence as a similarity metric to provide strong resistance to outliers and noise, and (ii) a self-updating process that supports the formation of singleton or extremely small clusters, enabling the generation of a large number of clusters when needed. Of note, use of γ-SUP clustering naturally circumvented the issues stemming from use of K-means as the clustering algorithm [1].

## Section 1. Development of a Robust and Efficient Clustering Method

### 1.1 Strengths and drawbacks of the original γ-SUP clustering algorithm

In developing RE2DC (Algorithm 1), we chose γ-SUP as its core clustering algorithm. Compared to existing methods for clustering particle images, γ-SUP uses (i) a self-updating process to support cluster formation initialized with individual data point as a singleton; (ii) q-Gaussian distribution to define a steep rejection boundary (q <1); (iii) γ-divergence as a similarity metric to provide strong resistance to outliers and noise. As a result, γ-SUP allows for formation of extremely small clusters, with promise of generating a large number of homogeneous classes from a massive and heterogeneous dataset, and naturally circumvents the issues of random initialization associated with use of K-means clustering [1]. However, the original γ-SUP has a number of practical shortcomings—its computation becomes costly when handling a large dataset (e.g., N > 10⁴ particles). In addition, γ-SUP needs selection of a hyper-parameter (see Algorithm 2). This hyper-parameter τ determines the number of clusters and the route of data points approaching their destiny clusters. Namely, when τ is extremely small, each data point forms its own cluster or a singleton, and as τ increases data points begin to merge into clusters. The selection of a good hyper-parameter for γ-SUP requires manual tuning, and it is determined based on inspecting a plot to identify a transition point (see Fig. 5 in [1]). In reality, the phase transition behavior might not always be evident. As a result, the selection of hyper-parameters τ is not only manual, but also subjective, where this hyper-parameter is often dataset-specific, all together making γ-SUP very difficult to use. To overcome these limitations, we launched algorithmic development with aim of (i) reducing computational complexity (CC); and (ii) eliminating the need for manual tuning of the hyper-parameter.

#### Algorithm 1 RE2DC

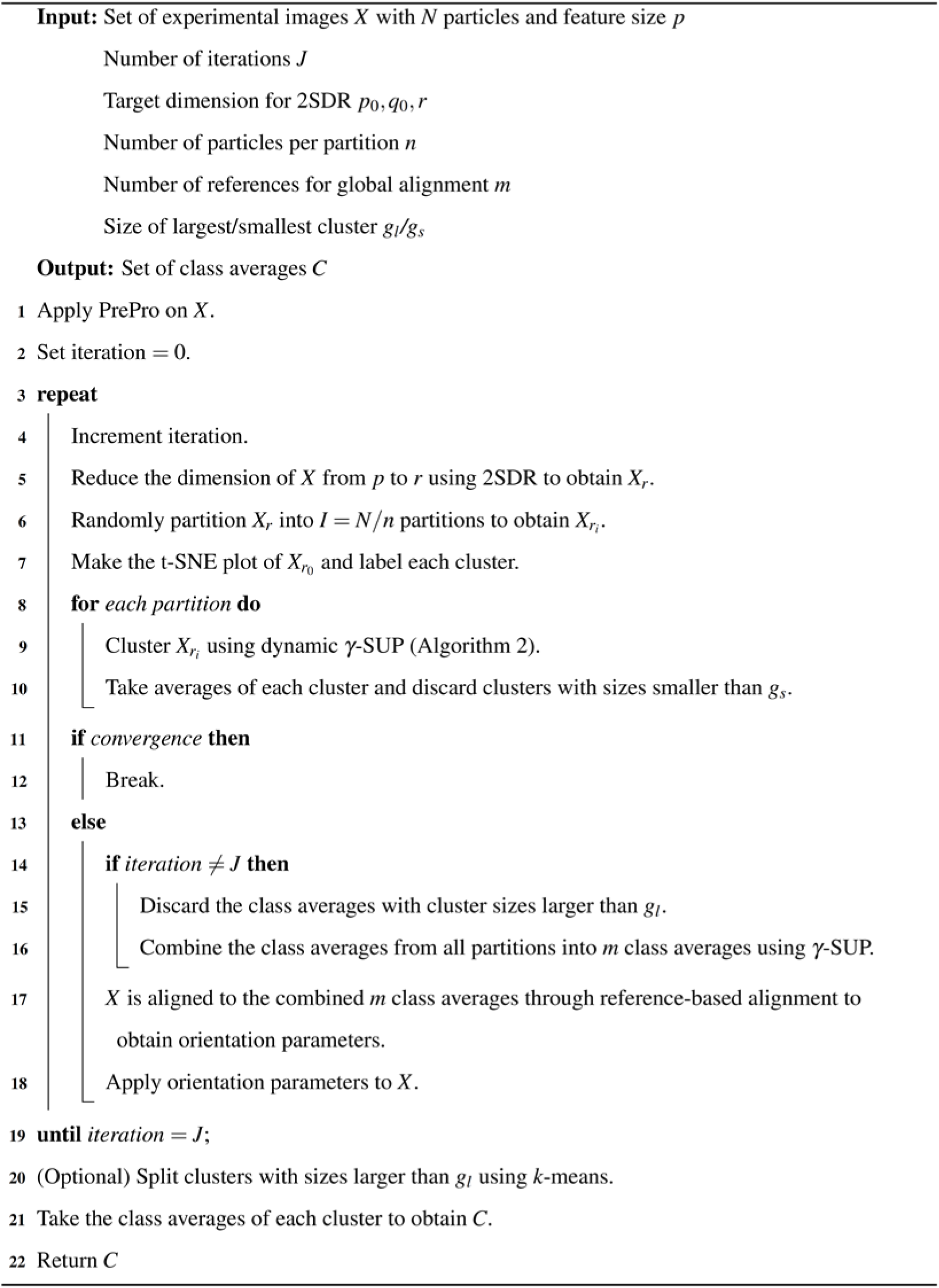

#### Algorithm 2 Dynamic γ-Sup

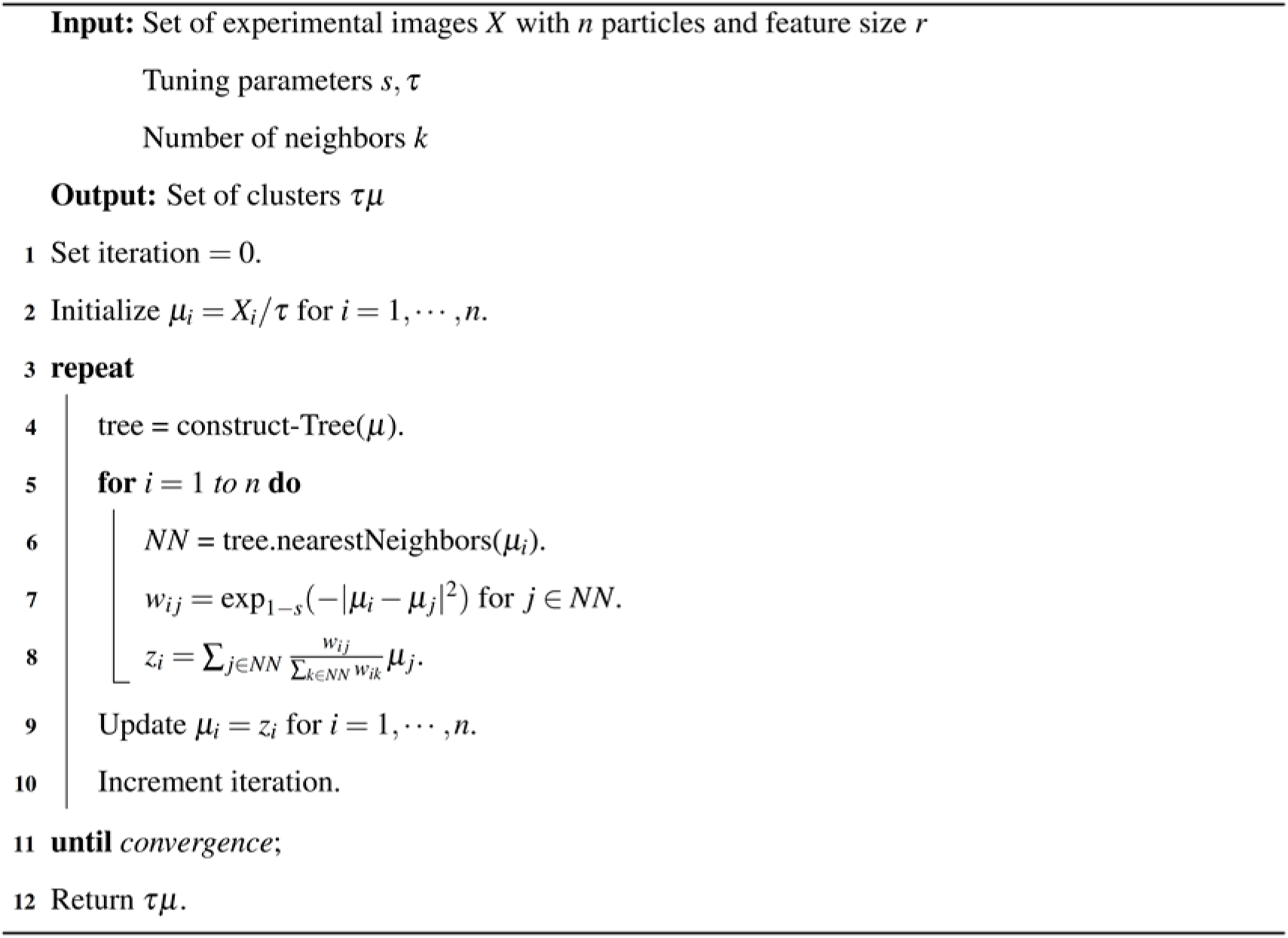

### 1.2 Reduction of γ-SUP computation complexity

We noticed that the weight matrix in γ-SUP is highly sparse, an inherent property of γ-SUP rooted in the use of γ-divergence as the metric. This property allowed one to use a sparse approximation for the weight matrix. To do so, we employed a well-known tree-based method [2] and constructed a k-nearest neighbor graph. To enable robust and efficient searching of the k nearest neighbors (kNN) for the sample points in each iteration, a space-partition tree was built and hierarchical graphs used [3]. The overall computation complexity (CC) of this kNN γ-SUP is thereby reduced from O(N^2^), that of original γ-SUP, to O(N log N), or even to O(N). In kNN γ-SUP, the number of k is fixed as 50 in accordance to the range of [5,150] suggested by the tree-based algorithm [2]. To further enlarge the range of search over iterations, we let k to grow with the number of iteration by setting it to be 10 + *j* ∗ 5 at iteration *j*—this dynamic adjustment strategy was inspired by a self-updating algorithm [4]. We named it “dynamic kNN γ-SUP” and further abbreviated it “dynamic γ-SUP”. Notably, this dynamic γ-SUP is highly efficient, empowered by a stage-wise data sampling strategy: (i) initial search that focuses on the local data structure, followed by (ii) expanded search by progressively increasing the number of neighbors.

### 1.3 Automating the determination of hyper-parameter in γ-SUP

To automate the selection of an optimal hyper-parameter, a two-stage procedure for its search was devised. In the first stage, a binary search is performed to identify a “non-trivial” region, a region that neither groups all sample points into a single cluster, nor assigns each sample point as a singleton. Next, exhaustive grid search is used on this region to find a specific value that maximizes a target function—e.g., particle yield, number of reasonable-sized clusters, or the combination.

### 1.4 Testing the efficiency of γ-SUP clustering algorithms using synthetic data

We then used numerical experiments to compare advanced versions of γ-SUP with original γ-SUP. To test their performance against ground truths, synthetic image data were used. They were generated as follows.

Using 5,000 experimental particle images of 70S ribosome handpicked by Haixiao Gao and Joachim Frank, we reconstructed a 3D structure using cryoSPARC [5]. We next generated 50 distinct 2D projection views from the 3D structure by projecting it in orientations equidistantly spaced in angular space. The projections were framed in a box of 130*×* 130 pixels with 2.82 Å pixel resolution. To create a balanced dataset, 5,000 images were produced from these 50 views by making 100 copies for each projection. Each projection’s 100 copies, from *i* = 1 to *i* = 100, underwent convolution with the electron microscope contrast transfer function based on the following parameters: 300 kV accelerating voltage, spherical aberration Cs = 2.0 mm, amplitude contrast = 0.07, and defocus values randomly selected from a set of 50 different values chosen from the range of 1.5 to 2 μm, while the astigmatism angle selected from the range of 0.2 to 1.4 radians. Independent and identically distributed Gaussian noise, represented as *N*(0,σ^2^), was added to each image. Three datasets with different signal-to-noise ratios (SNRs) 0.09, 0.06, and 0.03, were therefore produced.

An “unbalanced dataset” was also generated to emulate real data that may have preferred orientations. To do so, 10 views were chosen and replicated 400 times to present highly populated orientations, while the remaining 40 views were duplicated only 25 times to present less populated orientations.

For the scaling experiments described in Figure 2(c), datasets of increasing size (up to 50,000 images) were generated from the same 50 projection views by proportionally increasing the number of copies per view, preserving the unbalanced class-size ratio and SNR = 0.06.

## Section 2. Dimension Reduction Multi-reference Alignment (DRMRA)

### 2.1 Use of 2SDR to achieve dimension reduction and de-noising

Reliable and meaningful image clustering entails images to be first well aligned. However, in the presence of high noise, image alignment is highly error-prone where alignment solutions based on the highest cross-correlation scores do not warrant accurate or correct solutions. This issue of noise-vulnerability can, however, be mitigated by using de-noised particle images for alignment [6,7]. To do so, we used 2SDR, a tensor-based dimension reduction method [8], to denoise the images. As 2SDR is extremely rapid over related methods [6,7], it can be iteratively used along with every RE2DC iteration without causing noticeable time lag. With the use of 2SDR, we built a unique multi-reference alignment (MRA), termed dimension reduction multi-reference alignment (DRMRA). We then used DRMRA in each iteration of RE2DC (see Algorithm 1).

### 2.2 Use of Pre-Pro to initiate RE2DC

In addition to using 2SDR per RE2DC iteration, we initiated RE2DC using **Pre-Pro** [9] (see Algorithm 1). Notably, this **Pre-Pro** preprocessing incurs minimal computational overhead, and it has been used to initiate RELION or ISAC, significantly reducing the number of iterations required for converging to stable solutions.

### 2.3 Further acceleration of DRMRA by GPU

This DRMRA framework readily lends itself to GPU parallelization. To implement the GPU-acceleration, we used Cryo-RALib [10].

## Section 3. Use of Random Partition Strategy for Processing Massive Datasets

### 3.1 Random partition

To facilitate the processing of a massive dataset, we introduced a distributed computation strategy enabled by random partition of a large dataset into multiple small datasets. The distributed computation allows each small dataset to separately undergo 2D classification in a parallel manner. Random partition is offered as an option in RE2DC and it can be turned on when facing a large dataset (see Algorithm 1). Notably, this distributed computation strategy can be implemented on standard CPU hardware without requiring expensive hardware as does other parallel classification methods [11].

### 3.2 Partition parameter settings

To implement the partition, we recommend to set *n*, the number of particles per partition, to be 5000, and the number of references, *m*, to be 20. As for the upper and lower bounds of group size, *g_l_* and *g_s_*, 1000 and 15 are recommended respectively.

## Section 4. Implementation of RE2DC

RE2DC consists of two sub-algorithms, one for image alignment and the other for image clustering, as shown in Algorithm 1 and Algorithm 2. To initiate RE2DC, we employed Pre-Pro [9], which internally uses 2SDR to denoise particles, thereby facilitating improved initial alignment and accelerating convergence. As a result, most datasets typically require only three iterations to reach convergence. After initiation, 2SDR is further employed as a dimensionality reduction method in place of traditional PCA. This choice has been shown to offer computational advantages and is better suited for tensor-structured data such as cryo-EM images [8].

As for the sub-algorithm of image clustering (Algorithm 2), we used γ-SUP instead of K-means for clustering to avoid the shortcoming of existing classification methods largely rooted in K-means. In this development, we implemented and tested six versions of γ–SUP. These six versions are divided into two sets for the test. The first set includes the original one [1], a new one with k-nearest neighbors (kNN), which we term kNN γ–SUP, and a variant of kNN γ–SUP where k increases in each iteration, which we call ‘dynamic γ–SUP’ (Supplementary Figure 1). The three in the second set are variants of γ–estimator [1], which are the same as those in the first set, except that the sample points do not move in each iteration. The number of k is set to 50 in the kNN version and set to 10+ *j×* 5 in the dynamic version at iteration *j*. The clustering performance of the advanced variants of γ -SUP surpasses the original one in greatly reducing the computation time, as shown in Supplementary Figure 1. To combine DRMRA and the “dynamic γ–SUP” iteratively, we opted to minimize the number of initial references used to guide the MRA process. This way, the class assignment can be largely dictated by the clustering step. The class averages generated by the clustering step are further grouped by the dynamic γ-SUP into a smaller set of class averages as new references to guide the next round of image alignment. Furthermore, the class average that contains more than *g_l_* or smaller than *g_s_* will be discarded in the alignment step. This also has the benefit of speeding up the alignment step.

## Section 5. Using t-SNE visualization to display 2D classification result

### 5.1 Implementation of t-SNE

Visualizing high-dimensional data by projecting the data points into low dimensions has become an important method in many scientific applications. Numerous statistical tools have been developed for this end. Those approaches include PCA, Multidimensional Scaling (MDS), t-Distributed Stochastic Neighbor Embedding (t-SNE) [12]. A t-SNE algorithm comprises two main stages. First, it transforms the similarity matrix into a probability distribution (e.g., Gaussian distribution and t-distribution) for both the input data of high dimension and the visualization in low dimension (usually 2 or 3). Secondly, it minimizes the Kullback–Leibler (KL) divergence between the two distributions by a gradient descent algorithm. In practice, under a suitable parameter setting, t-SNE can effectively preserve the high-level qualitative structures of the input data, such as the patterns of nearest clusters or groups [13]. As the history of data visualization methodology is concerned, t-SNE was the first successfully used to separate the famous MNIST dataset into 10 clusters. Since that success, t-SNE has been widely applied to exploring the heterogeneity of high dimensional data with applications in diverse areas [13,14], including cancer biology and bioinformatics.

To date, the process of 2D classification of cryo-EM particle images is virtually treated as a black box for an end-user, and it is therefore difficult to have an intuitive insight into the progress or the quality of classification results. Here, to unveil this black box, we introduced t-SNE, enabling projection of data in high-dimension space onto low dimensions for visualization.

In practice, use of t-SNE usually requires PCA preprocessing for reducing the dimensionality of the data. This incurs cumbersome overhead when facing cryo-EM images with large number of pixels (>10**⁴**). To circumvent this, we replaced those PCA steps with 2SDR. In addition, we employed a fast t-SNE that is faithful in preserving global data structure. This is based on FFT-accelerated interpolation [14] and initialization scheme using scaled PCA [13].

While we were applying t-SNE to visualize 2D classification results of 70S ribosome, we surprisingly found that in some of the data points that were “considered” by the classification algorithm to belong to the same group can be scattered across the t-SNE plot, indicating that those data points actually belong to different clusters but were “forcefully” grouped together by the algorithm being tested, as shown by the class of the central panel in Supplementary Figure 5. As a result, the corresponding class average is fairly blurred. In comparison, in the class shown in the right panel of Supplementary Figure 5, the red points assigned by the algorithm are localized within the same cluster. In this case, the corresponding class average exhibits clearer structural details. This discovery suggests that t-SNE plot could be used to evaluate the quality of the classification results or a classification algorithm.

### 5.2 Interactive t-SNE

To allow a user to directly interact with the t-SNE plot to refine classification results, we employed holoviz/hvplot [15], a high-level plotting API, to build an interactive interface for interactive particle selection and filtering. The interface allows the user to inspect each part of the t-SNE landscape to fine-tune the classification results by choosing a region for averaging (Figure 3(d)), which is feasible even when the particles are crowded illustrated in Supplementary Figure 6.

## Section 6. Data description and preparation

Prior to running RE2DC for 2D classification, users can opt to correct contrast transfer function (CTF). The CTF-estimation could be performed at the micrograph level or at individual particle level prior to particle extraction. When using RE2DC on ASCEP, users can apply “phase flipping” to particle images for CTF correction, as commonly undertaken by popular classification methods. When handling images with large size, down-sampling of images is recommended. This strategy accompanied by applying a circular mask to each particle image is commonly used by many algorithms. Although down-sampling helps minimize image noise, it removes high-resolution details in images that the resulting 2D class averages can appear coarse. However, when retaining fine structure details in 2D class averages is demanded, RE2DC offers an option that allows for generating class averages using original images while leveraging pose parameters inferred based on down-sampled images.

In this study, six experimental cryo-EM datasets are used for the tests. All the data we used has been down-sampling by about 5X to speed up the tests.

The first set is E. coli 70S ribosome recorded in Joachim Frank lab using a CCD camera. The original set contains 10,000 boxed-out particles with a box size of 130*×* 130 pixels of 2.82 Å and is a widely used test dataset. We only used the first half set, which is ribosome bound with an elongation factor (EF-G). We termed it 70S ribosome dataset.

The second set is the GS protein particles we collected using a Titan Krios operated at 300 kV and a Falcon III direct electron camera (ThermoFisher Scientific) hosted by Academia Sinica Cryo-EM Center (ASCEM). The total electron dose was 40 e^-^/Å ^2^ distributed in 10 movie frames. The movie data was motion-corrected using MotionCor2 [16], CTF-corrected using CTFFIND4 [17], with initial particle pick performed using CryoSPARC pipeline [5]. This set contains a total of 94,154 particles.

The third dataset is malaria 80S ribosome particles from Scheres lab [18]. The particle images were downloaded from [EMPIAR-10028], https://www.ebi.ac.uk/pdbe/emdb/empiar/entry/10028/. This set contains 105,247 particles carefully selected from a larger set of 158,212 particles by the authors and is provided as a RELION Benchmark example. This data was recorded on a direct camera (Falcon II) using a 300 kV Titan Krios cryo-EM at MRC. It is used to test the performance of a cryo-EM image processing algorithm. The size of the particle image is 360*×* 360 pixels of 1.34 Å. The defocus parameters were provided for each particle by the authors.

The fourth dataset is TRPV1 particles downloaded from [EMPIAR-10005], https://www.ebi.ac.uk/pdbe/emdb/empiar/entry/10005/. The original dataset was recorded on a direct electron counting camera (K2) with super-resolution mode (0.6 Å) in Yifan Cheng lab at UCSF using a 300 kV cryo-EM with a side-entry holder [19] and then extracted and decimated by a factor of 2 into images with a box size of 256*×* 256 pixels (1.2 Å) by the authors. The dataset contains 88,915 particles. We term it “NC-TRPV1”.

The fifth dataset is TRPV1 particles embedded in nanodisc, downloaded from [EMPIAR-10059], https://www.ebi.ac.uk/pdbe/emdb/empiar/entry/10059/. The dataset is with a box size of 192*×* 192 pixels (1.2 Å) by the authors [20]. All 218,805 particles were used in this study. We term it “NanoD-TRPV1”.

The last one is movies of SARS-CoV-2 Spike stabilized in prefusion state, from [EMPIAR-10514], https://www.ebi.ac.uk/pdbe/emdb/empiar/entry/10514/ [21]. The dataset contains 3,511 movies with pixel size 1.08 Å. To deblur the micrographs, we performed motion correction using the movie frame alignment function with MotionCor2 [16]. The contrast transfer function (CTF) estimation was performed by CTFFIND4 [17], and the correction was applied to each micrograph. Finally, using crYOLO [22] auto-picking function, we extracted 375,067 particles from these micrographs with a box size of 400*×* 400 pixels.

## Section 7. 3D reconstruction using CryoSPARC

3D reconstruction from clean particle sets obtained by RE2DC or other 2D classification methods was performed using CryoSPARC [5] with ab initio model followed by 3D non-uniform refinement [23]. The overall resolution was assessed based on the standard Fourier shell correlation (FSC) at 0.143 and reported using cryoSPARC [5].

## Section 8. Computation Resources

As for the computers used for the computation, all the experiments are run on a workstation equipped with two Intel Xeon CPU E5-2699 v4 at 2.20GHz and eight NVIDIA Geforce GTX 1080 Ti graphics cards.

## Supplementary Table

**Supplementary Table1:** Comparison of classification time and particle yield across six datasets.

| Algorithm | Dataset | Performance |  |  |
| --- | --- | --- | --- | --- |
|  |  | Execution Time<br>(CPU/GPU) (Hour)<sup>a</sup> | Number of Good<br>Classes | Particle Yield/<br>Resolution(A)@0.143 |
| RELION | 70S ribosome (30 classes) | 0.43/<u>0.28</u> | 15 | 4,240 |
|  | GS Protein (150 classes) | 22.91/<u>8.68</u> | 40 | 91,266 |
|  | 80S ribosome (520 classes) | 62.32(f)/<u>30.12(f)</u> | 141 | 102,419 |
|  | NC-TRPV1 (200 classes) | 16.03(f)/<u>13.56(f)</u> | 15 | 50,620/3.43 |
|  | NanoD-TRPV1 (100 classes) | 12.36(f)/<u>9.49(f)</u> | 20 | 153,839/2.96 |
|  | SARS-CoV-2 (500 classes) | 120.03(f)/<u>50.21(f)</u> | 10 | 235,300/3.48 |
| ISAC | 70S ribosome (200 members) | <u>1.78</u>/0.33 | 29 | 3,875 |
|  | GS Protein (1000 members) | <u>50.92</u>/4.08 | 98 | 87,665 |
|  | 80S ribosome<br>(4X binning, 200 members) | <u>124.50</u>/14.55 | 514 | 103,043 |
|  | NC-TRPV1<br>(3X binning, 1000 members) | <u>55.61</u>/5.38 | 55 | 50,798/3.39 |
|  | NanoD-TRPV1<br>(3X binning, 1000 members) | */<u>12.65</u> | 138 | 125,656/2.93 |
| RE2DC | 70S ribosome | 0.28/<u>0.18</u> | 27 | 4,667 |
|  | GS Protein | 1.82/<u>1.58</u> | 169 | 89,588 |
|  | 80S ribosome (4X binning) | 2.54/<u>2.02</u> | 542 | 101,863 |
|  | NC-TRPV1 (3X binning) | 1.76/<u>1.51</u> | 88 | 59,106/3.38 |
|  | NanoD-TRPV1 (3X binning) | 3.86/<u>3.33</u> | 317 | 159,988/2.91 |
|  | SARS-CoV-2 (5X binning) | 8.45/<u>7.76</u> | 442 | 221,916/3.44 |

The Fast-subset is utilized for RELION when feasible. For RE2DC, partition sizes are set as follows: 5,000 for 70S ribosome, 80S ribosome, and NC-TRPV1; 20,000 for SARS-CoV-2; and 10,000 for other datasets. Dynamic $\gamma$ -SUP is employed during the clustering phase. We utilize 44 Message Passing Interface (MPI) processes and a single GPU. The execution time represents an average of 5 replicas for the 70S ribosome and 3 replicas for all other datasets. Note that the data format conversion time on the platform is excluded from our count. The statistics for particle yield, the number of good classes, and the resolution of 3D reconstruction are indicated in the last two columns. An asterisk (\*) indicates that the run had failed.

## Supplementary Figures

**Supplementary Fig. 1.**
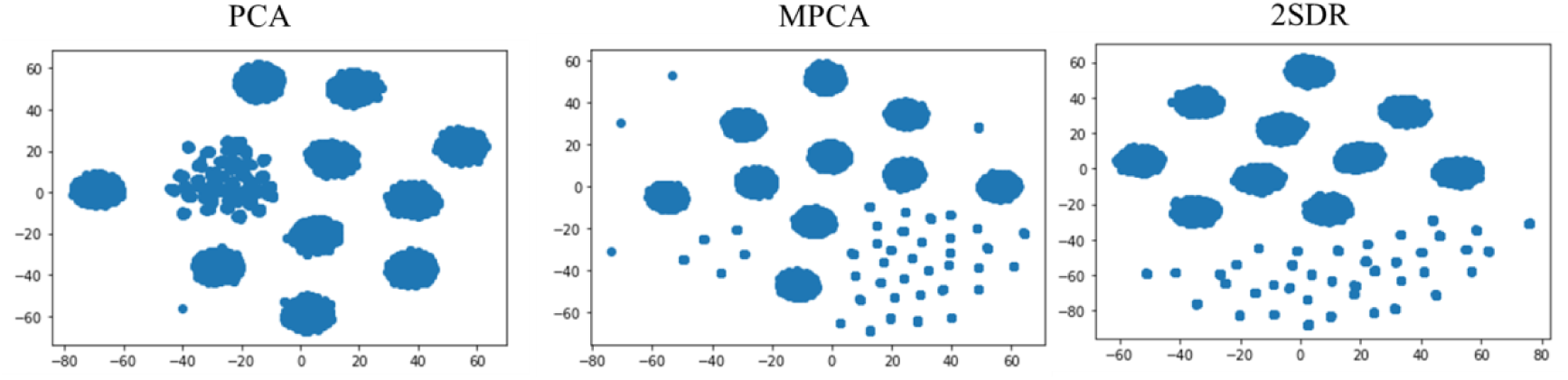
t-SNE visualization of the unbalanced set of synthetic data using different preprocessing tools for dimension reduction, from left to right: PCA, MPCA, and 2SDR.

**Supplementary Fig. 2.**
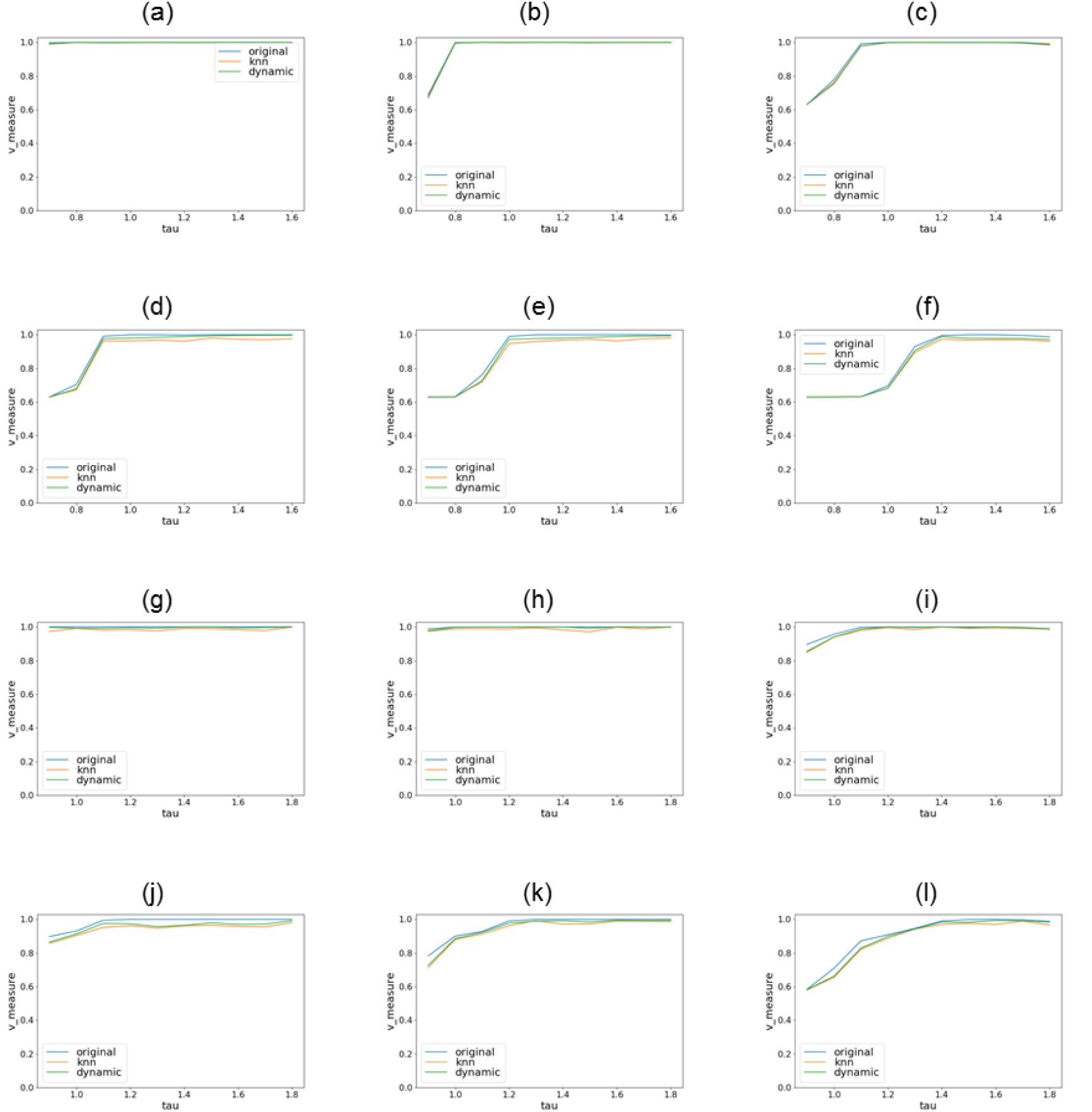
V-measure of the clustering accuracy for various γ-SUP variants. **(a)**, **(b)**, & **(c)**: the V-measure with respect to τ hyper-parameters for various γ-SUP variants on the balanced dataset with SNRs of 0.09, 0.06, and 0.03, respectively. **(d)**, **(e)**, & **(f)**: the V-measure with respect to τ hyper-parameters for various variants of γ-estimator, the predecessor of γ-SUP, on the balanced dataset with SNRs of 0.09, 0.06, and 0.03, respectively. **(g)**, **(h)**, & **(i)**, mirror **(a)**, **(b)**, & **(c)** for the unbalanced dataset. **(j)**, **(k)**, & **(l)**, mirror **(d)**, **(e)**, and **(f)** for the unbalanced dataset.

**Supplementary Fig. 3.**
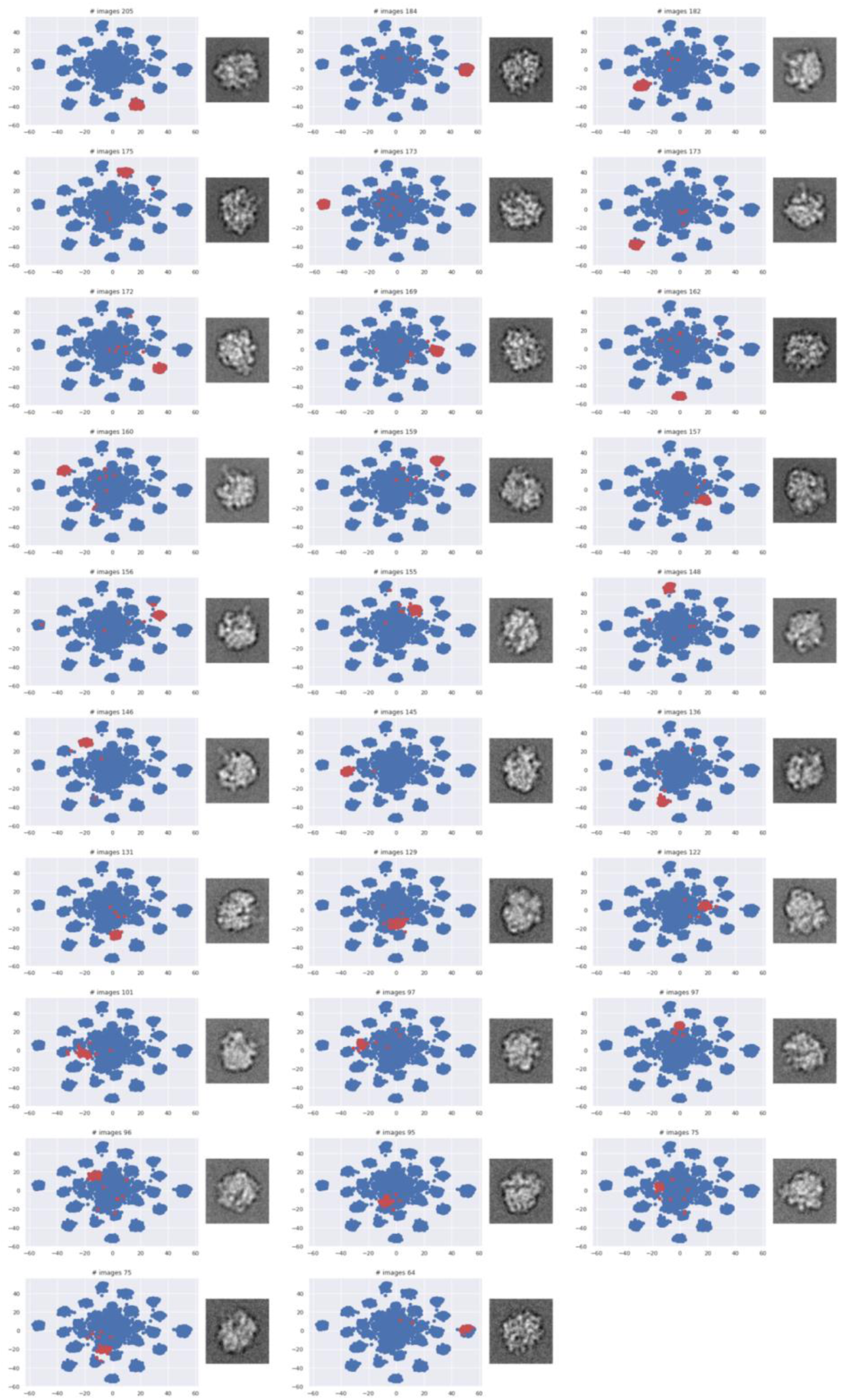
t-SNE sub-plots for the classification results of the 70S ribosome using ISAC, with 200 set as the class size limit. Blue dots represent data points grouped together by the alignment step in ISAC, while red dots represent data points assigned to the same class by the clustering step in ISAC. The image next to each t-SNE sub-plot is the corresponding class average, namely the averages from the red dots.

**Supplementary Fig. 4.**
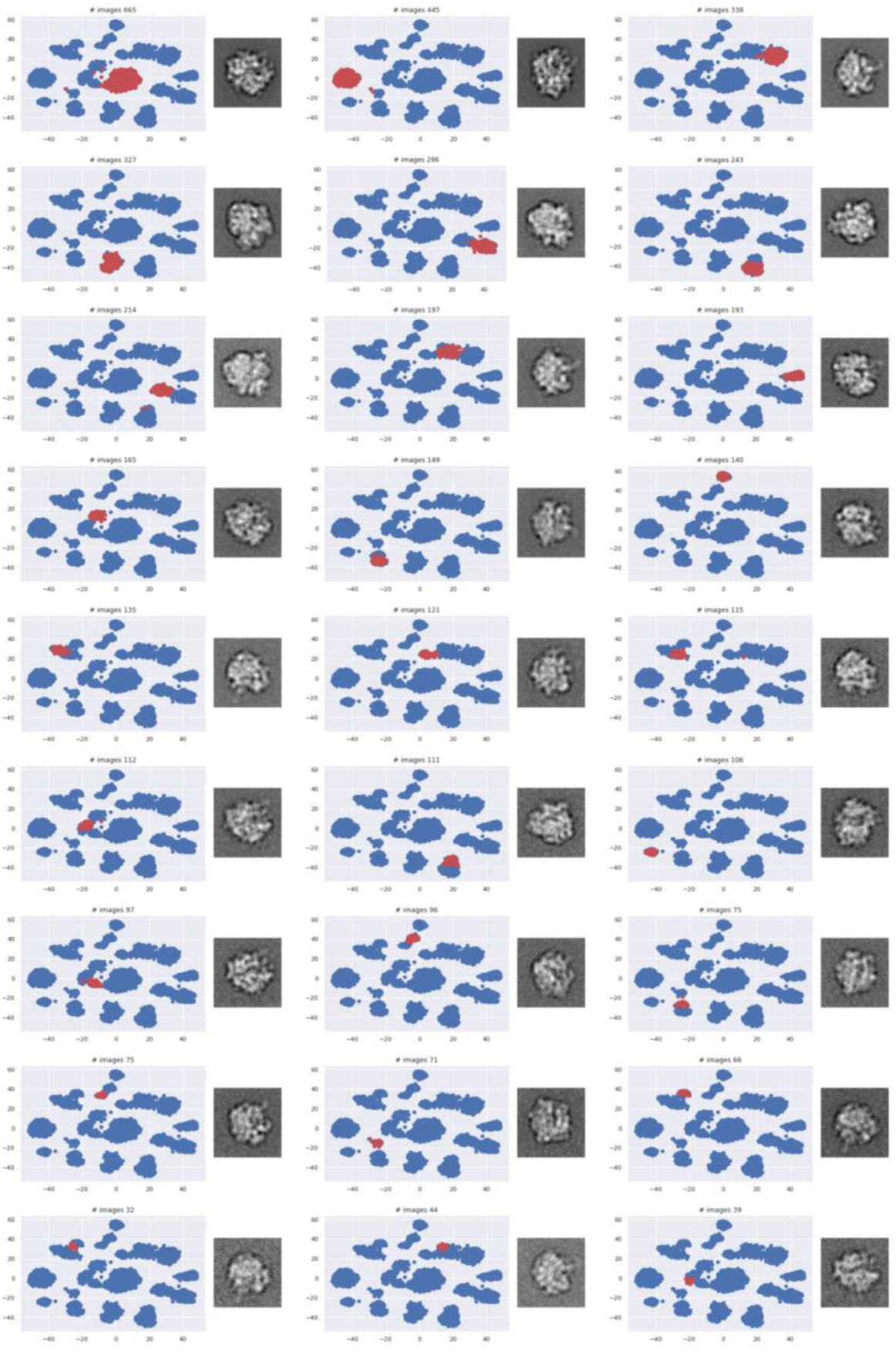
t-SNE sub-plots for the classification of the 70S ribosome using RE2DC for clusters with members more than 15. Blue dots represent data points grouped together by the alignment step in RE2DC, while red dots represent data points assigned to the same class by the clustering step in RE2DC. The image next to each t-SNE sub-plot is the corresponding class average, namely the averages from the red dots.

**Supplementary Fig. 5.**
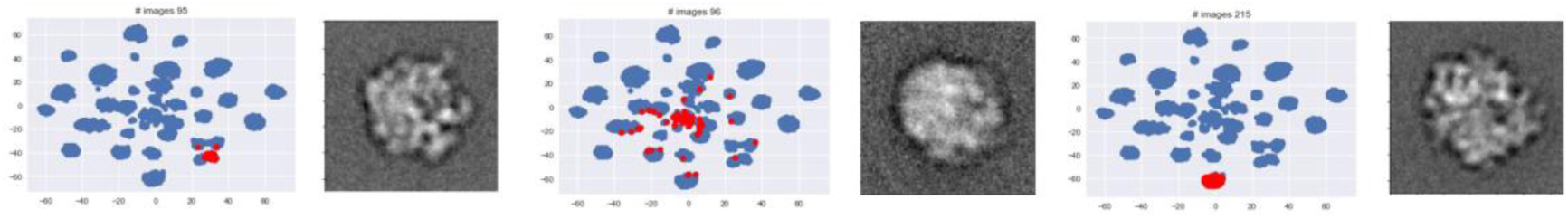
Three representative t-SNE sub-plots along with the class averages from a ISAC 2D classification experiment. The blurred class average in the middle does not indicate the members in this class are “bad particles”, they merely not “alike” particles but forcefully assigned into the same class by ISAC.

**Supplementary Fig. 6.**
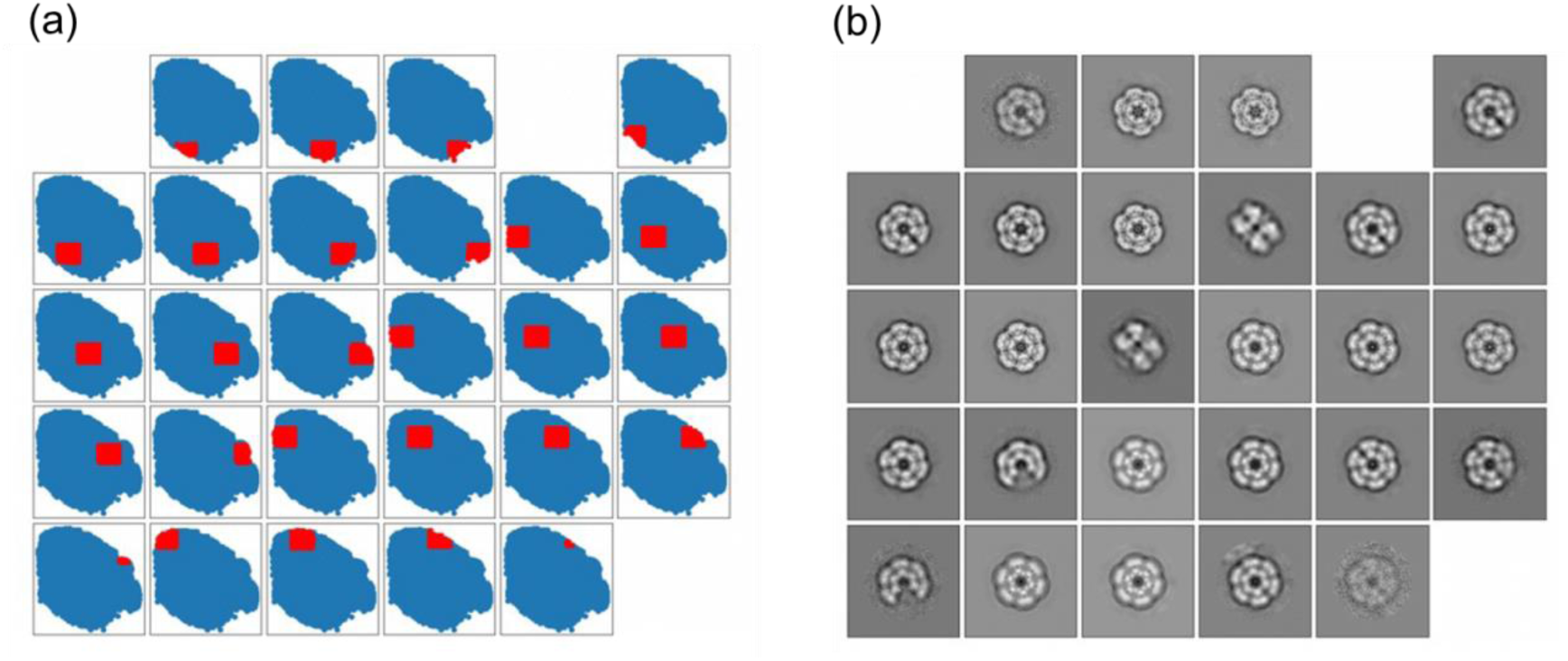
Use of i-t-SNE to produce classes along with class average images under crowding conditions. **(a)** Illustration of the grid-based approach when the data points are crowded, each grid colored in red. **(b)** The class averages from data points in each grid in **(a)**.

**Supplementary Fig. 7.**
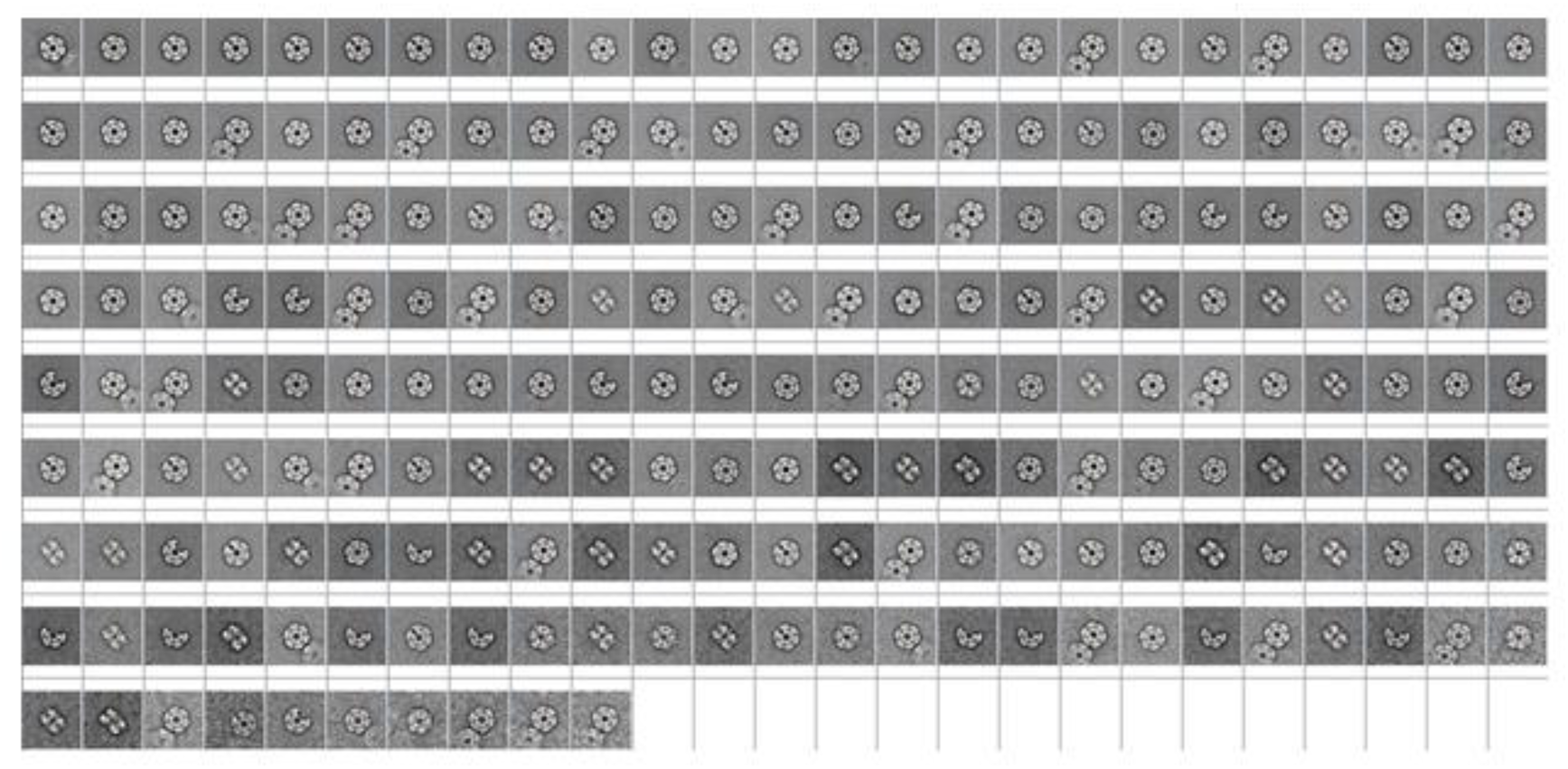
Classification results of GS protein using RE2DC.

**Supplementary Fig. 8.**
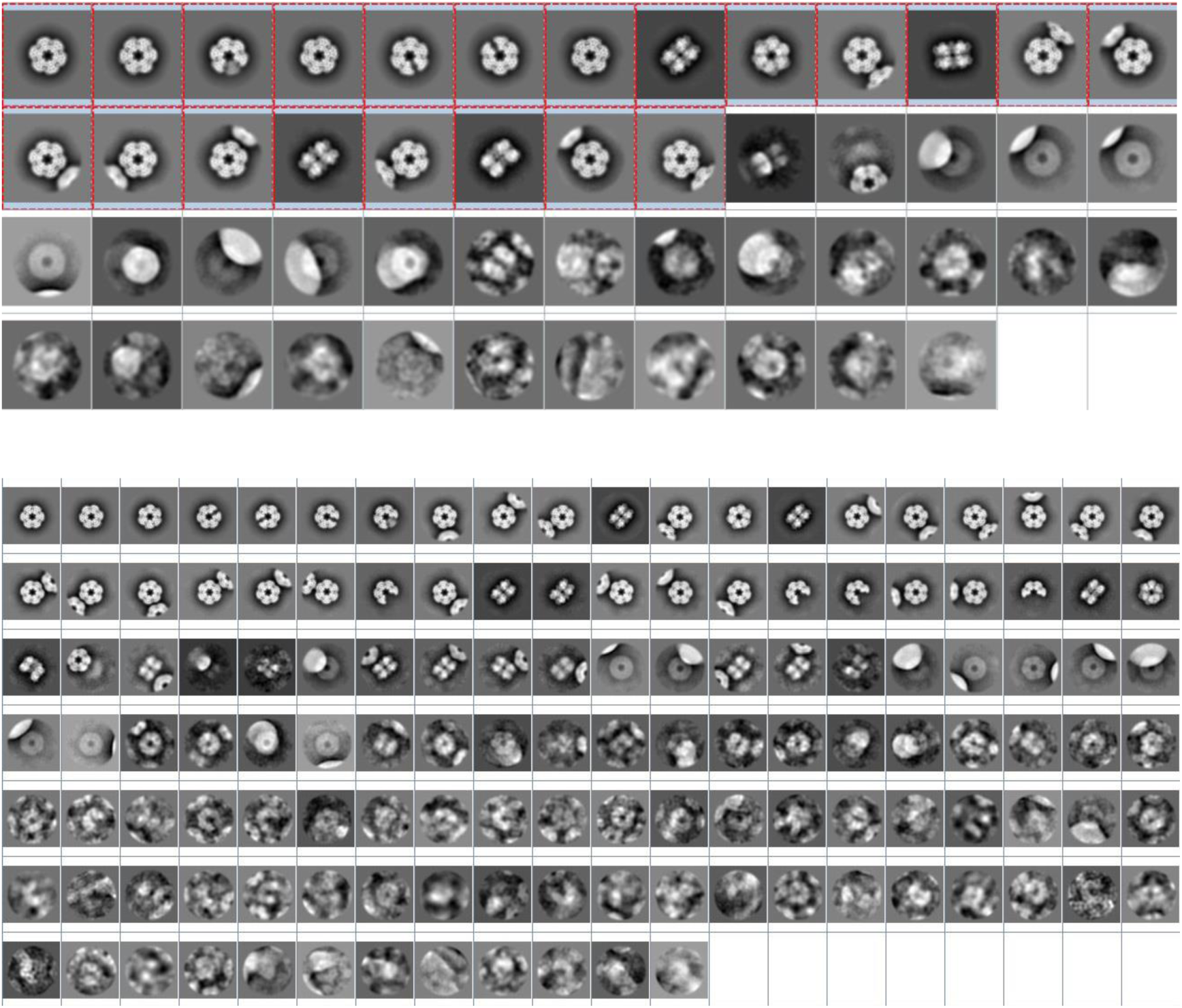
Classification results of GS protein using RELION. Upper Panel : RELION with 50 classes. Lower Panel: RELION with 100 classes.

**Supplementary Fig. 9.**
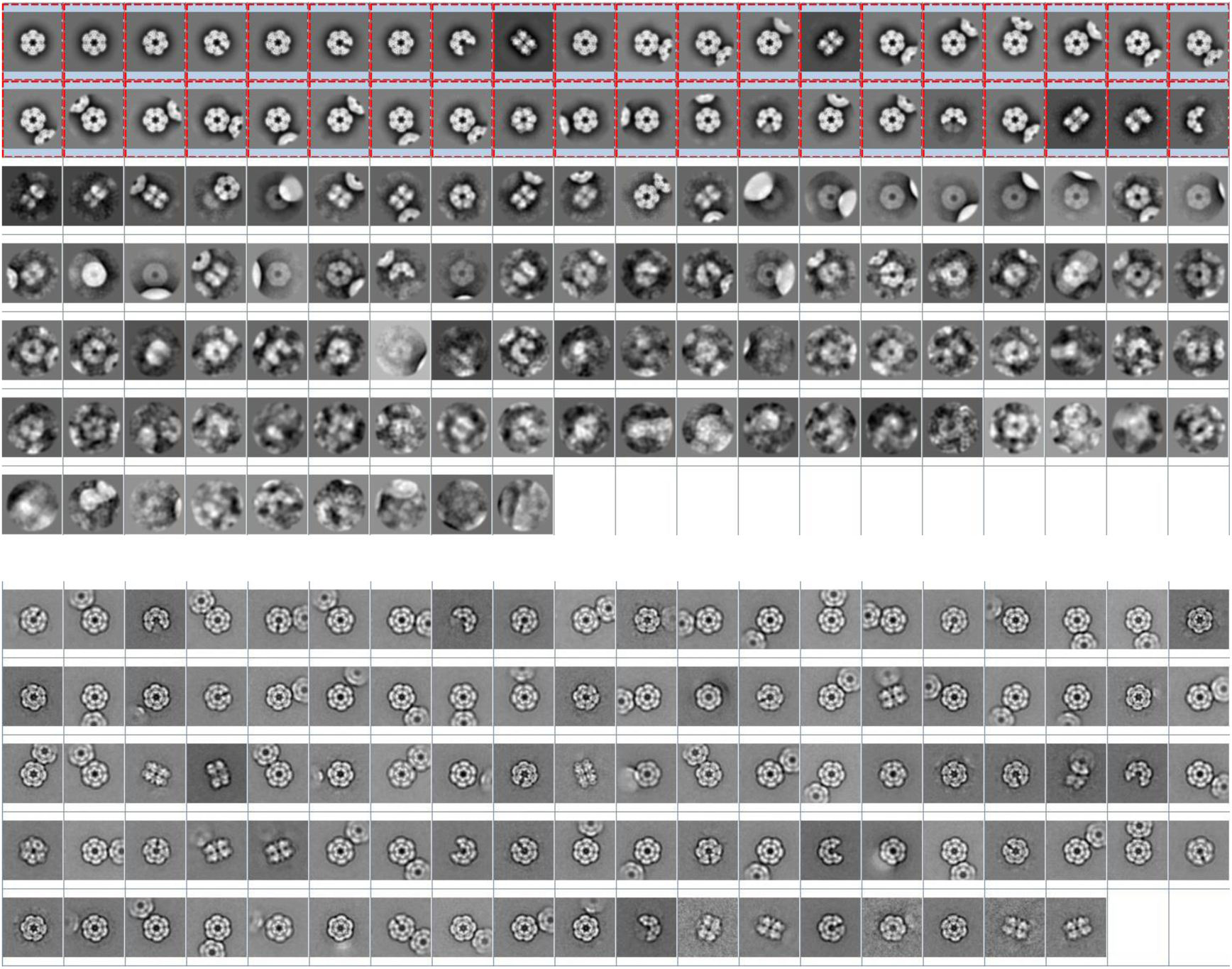
Classification results of GS protein using ISAC. Upper Panel: ISAC with 2000 as the size limit per class. Lower Panel: ISAC with 1000 as the size limit per class.

**Supplementary Fig. 10.**
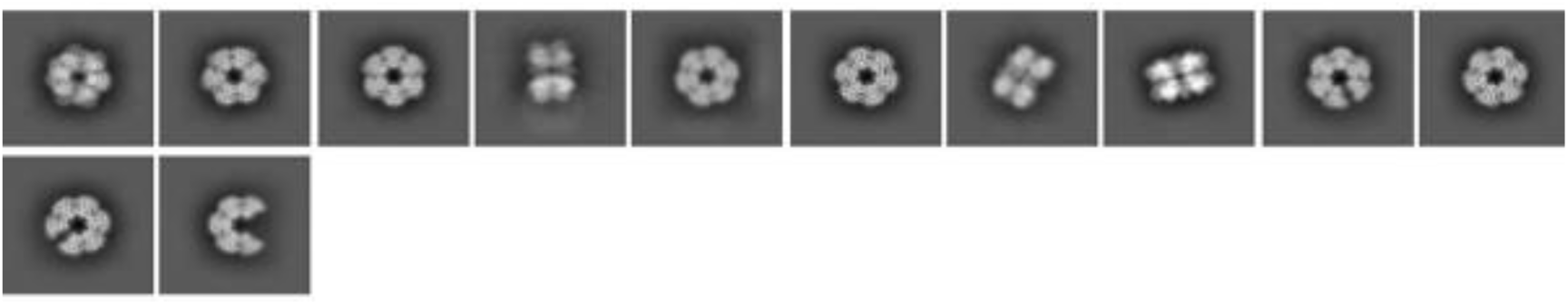
Classification results of GS protein using CryoSPARC.

**Supplementary Fig. 11.**
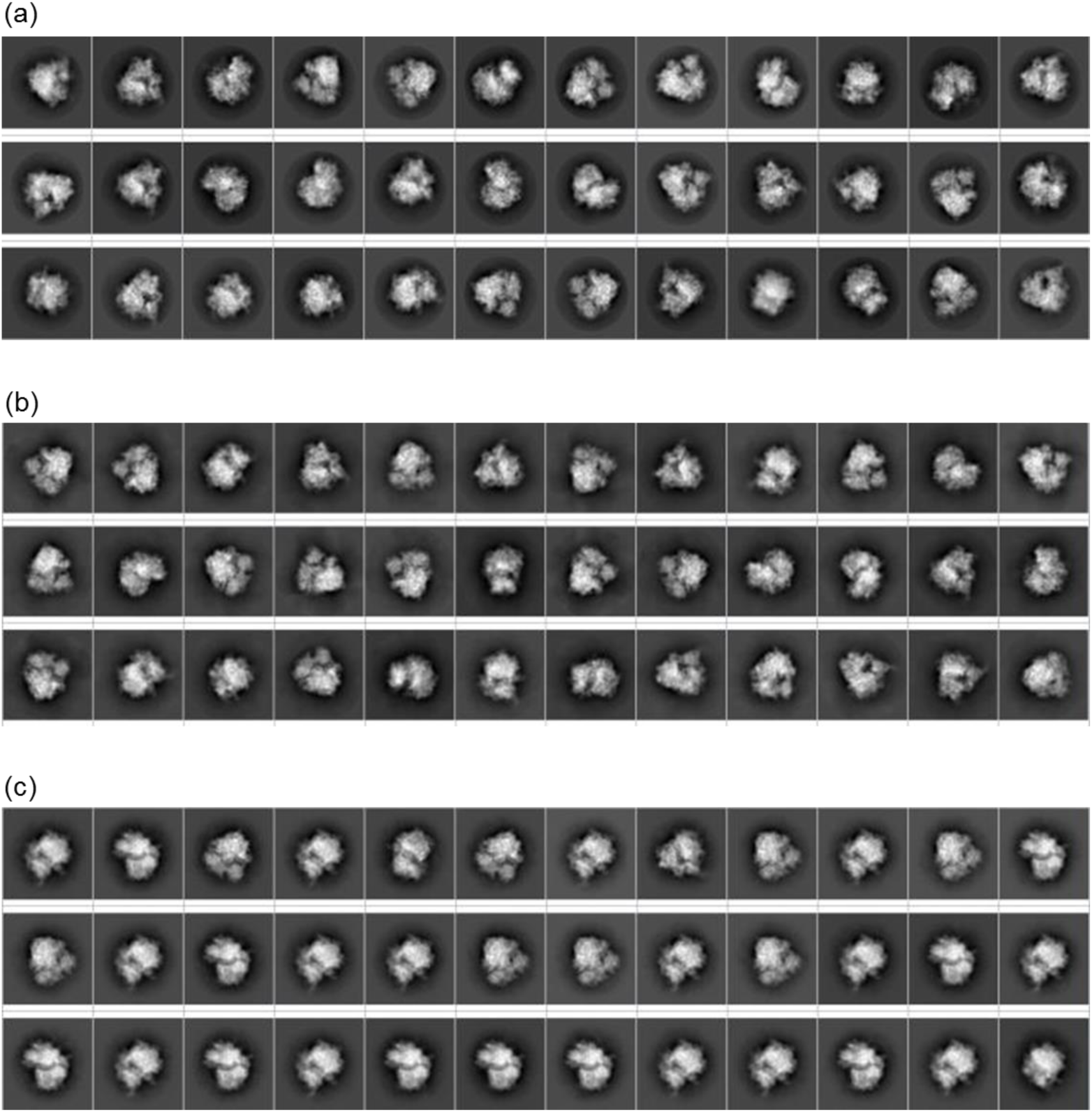
Representative class averages resulted from the 2D classification on an already curated 80S particle dataset [EMPIAR-10028] using three different methods. (a) RELION. (b) ISAC. (c) RE2DC.

**Supplementary Fig. 12.**
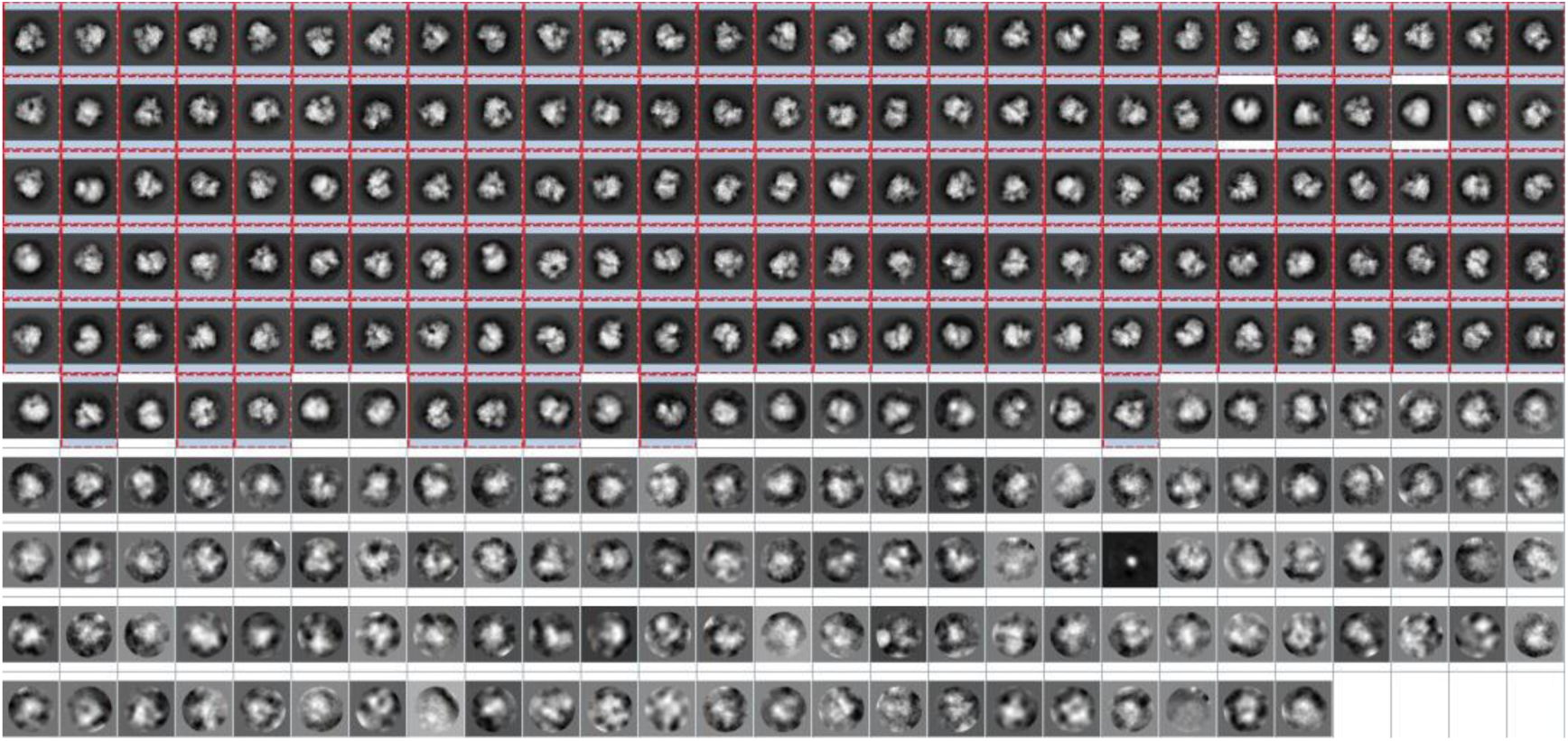
Complete classification results of 80S ribosome using RELION: (a) with the 520 set as the pre-specified class number. A total of 266 classes are produced where the 141 high-quality classes are boxed in red.

**Supplementary Fig. 13.**
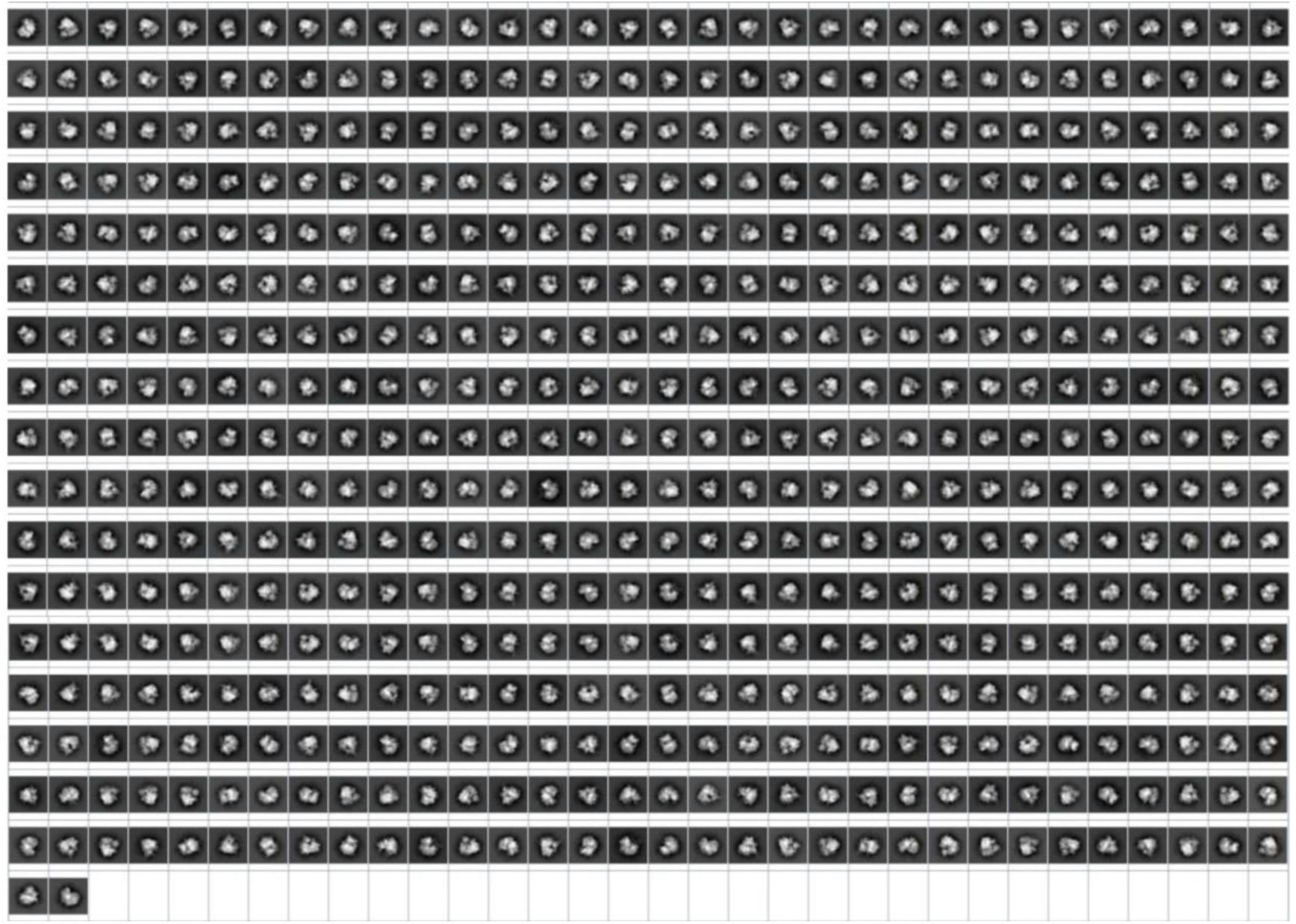
Complete classification results of 80S ribosome using ISAC with class size limit set to be 200.

**Supplementary Fig. 14.**
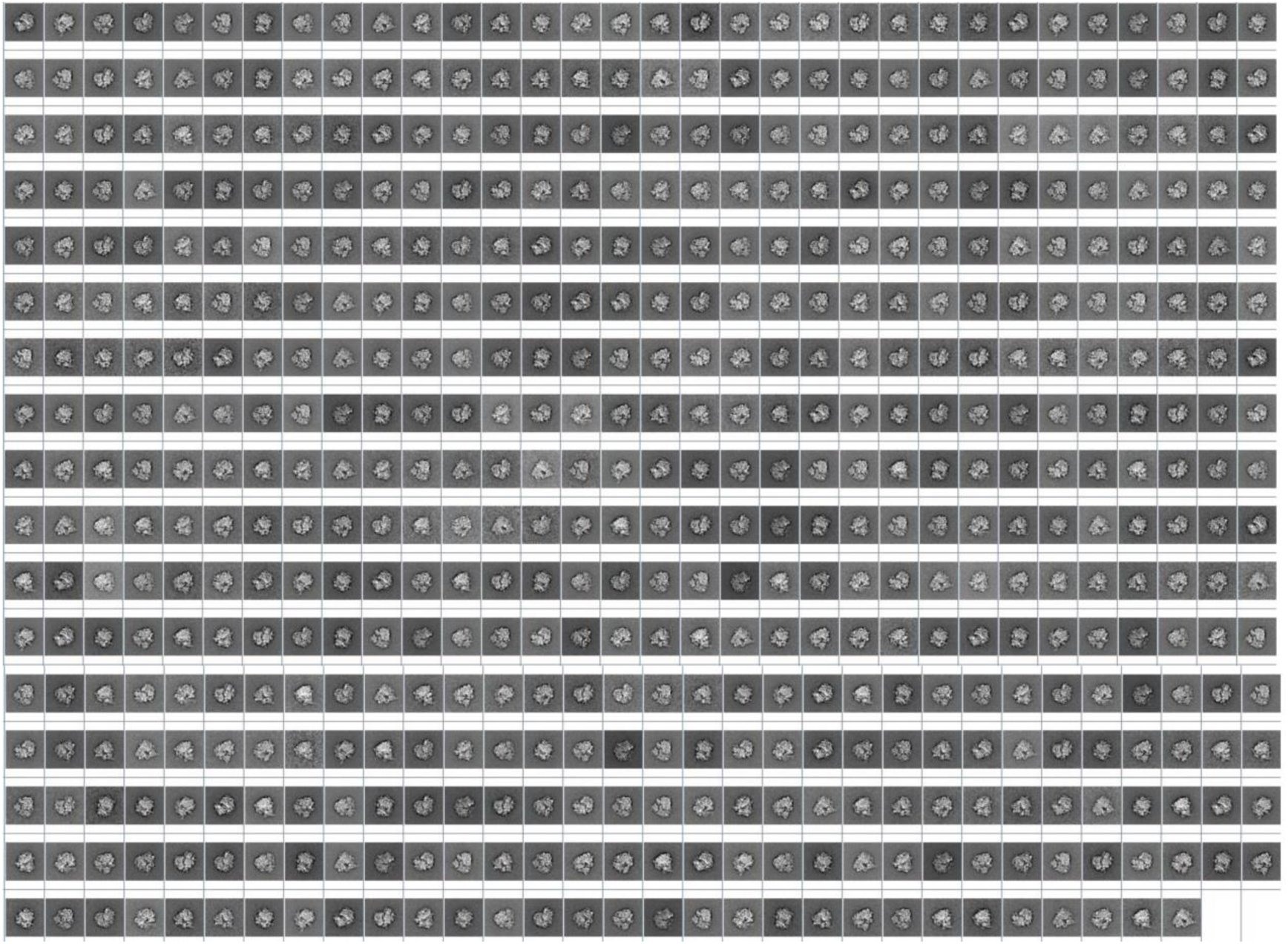
Complete classification results of 80S ribosome using RE2DC with the least number per class set to be 15.

**Supplementary Fig. 15.**
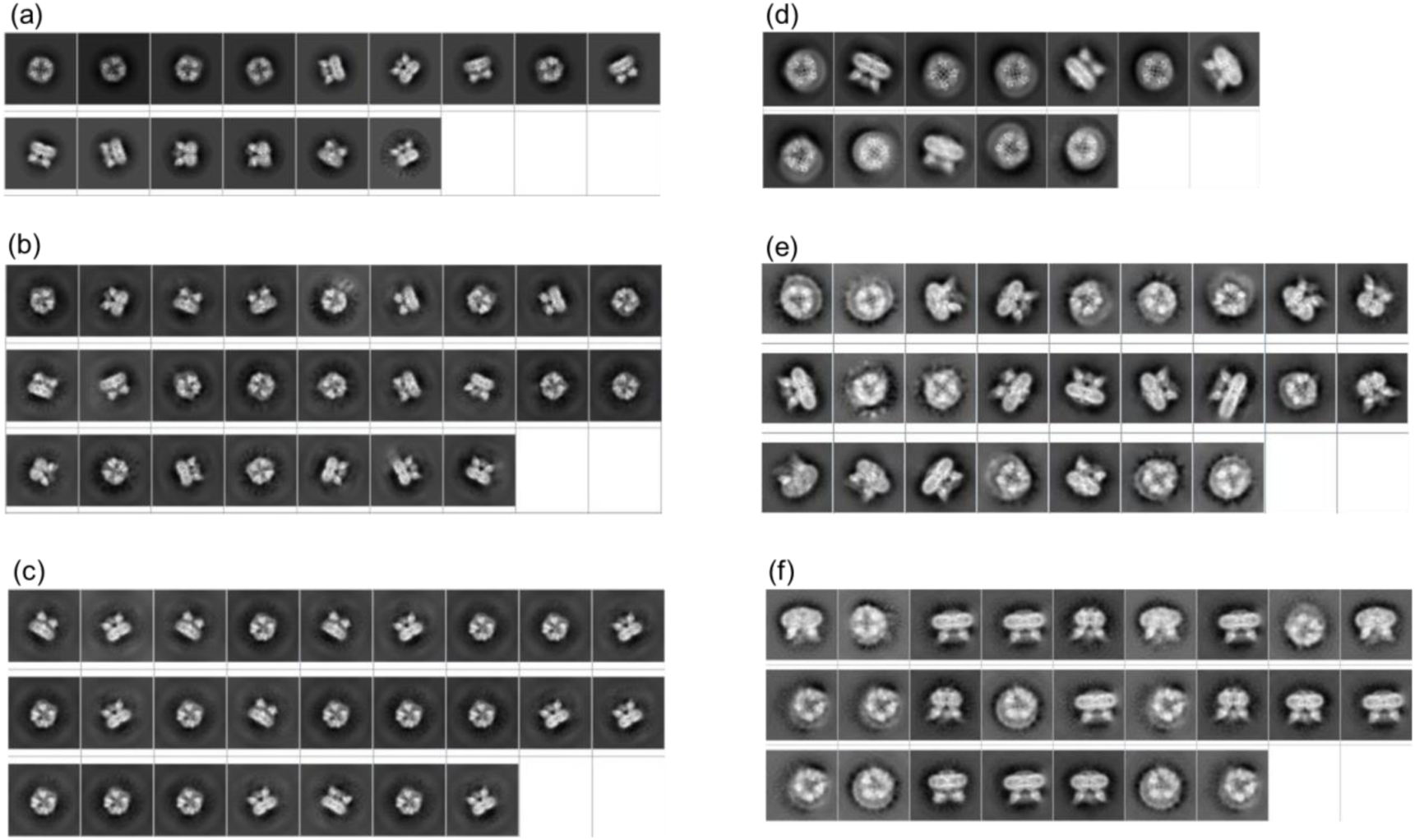
2D classification results for two non-curated TRPV1 channel datasets using three different classification methods. **(a)**-**(c)** NC-TRPV1 [EMPIAR-10005] using RELION, ISAC, and RE2DC respectively; **(d)**-**(f)** NanoD-TRPV1 [EMPIAR-10059] using RELION, ISAC, and RE2DC respectively.

**Supplementary Fig. 16.**
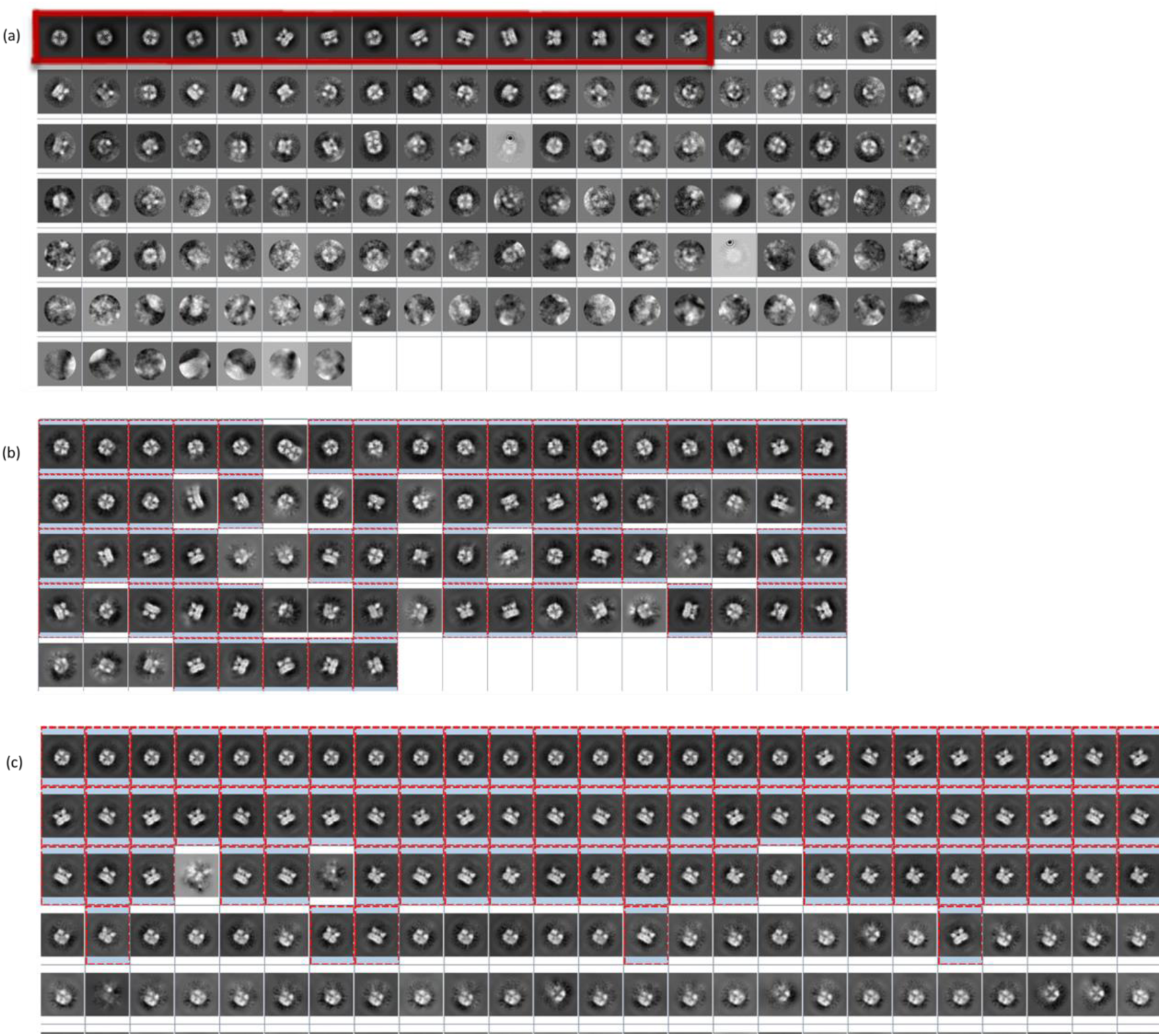
Overall classification results of NC-TRPV1 using three different classification methods. The high-quality class averages are boxed in red. (a) RELION with 200 as prescribed class number. (b) ISAC with class size limit set to 1000. (c) RE2DC.

**Supplementary Fig. 17.**
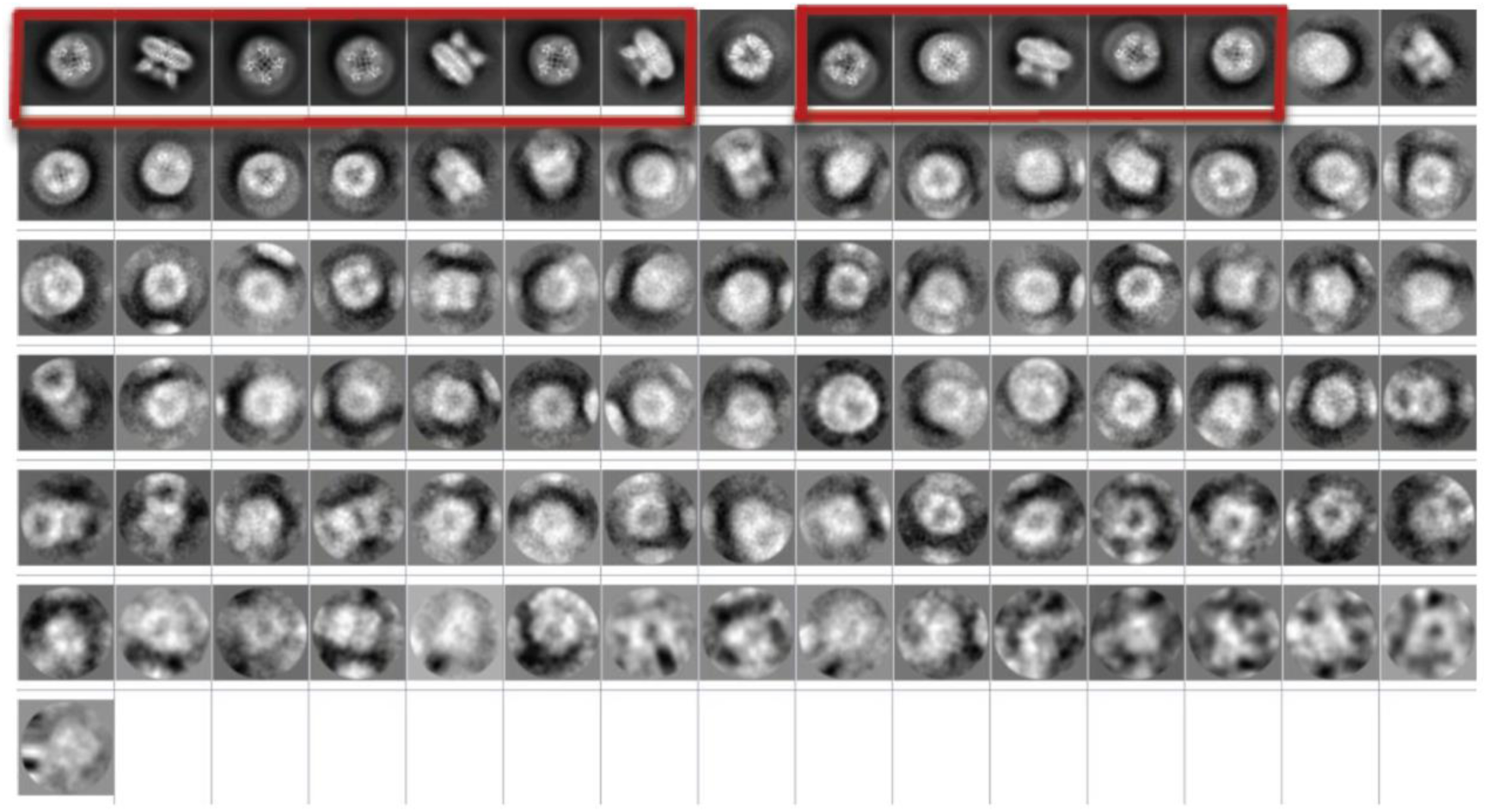
Classification results of NanoD-TRPV1 using RELION with 100 set as the pre-specified class number.

**Supplementary Fig. 18.**
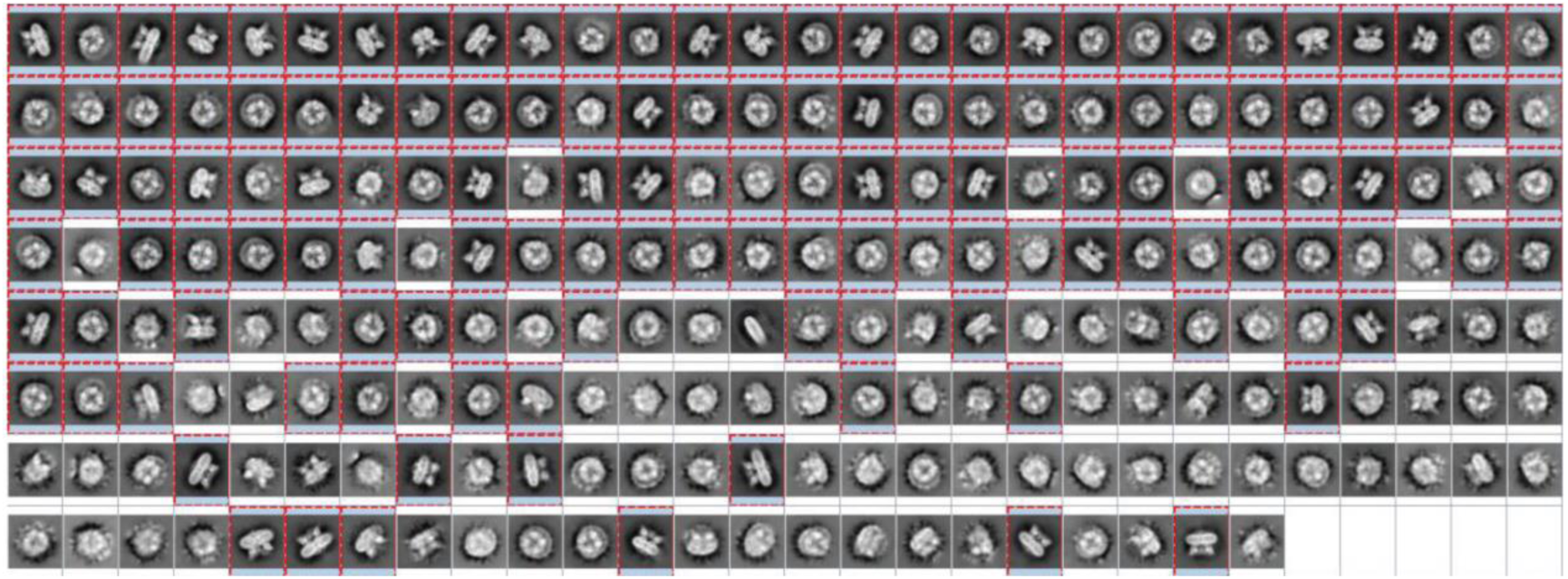
Classification results of NanoD-TRPV1 using GPU-ISAC with class size limit set to be 1000.

**Supplementary Fig. 19.**
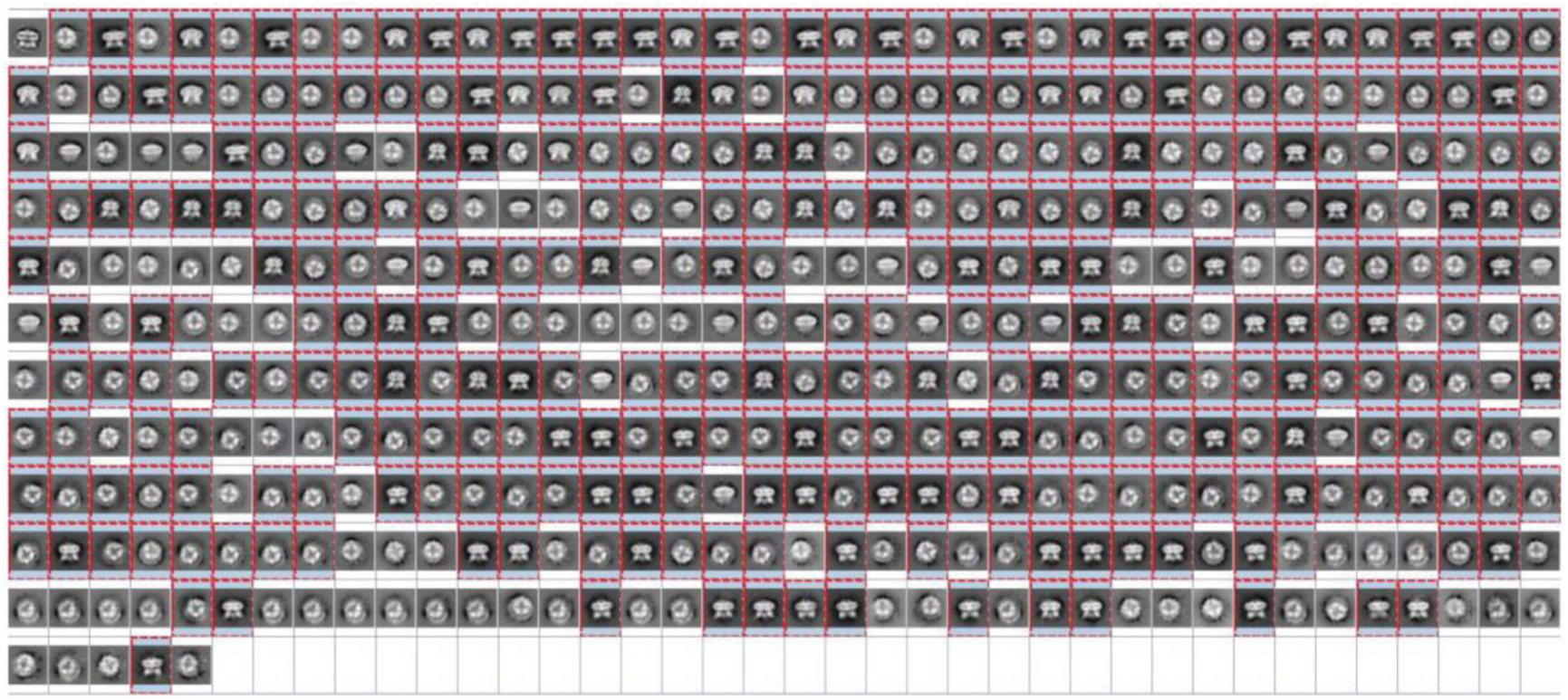
Classification results of NanoD-TRPV1 using RE2DC. The high-quality class averages are boxed in red.

**Supplementary Fig. 20.**
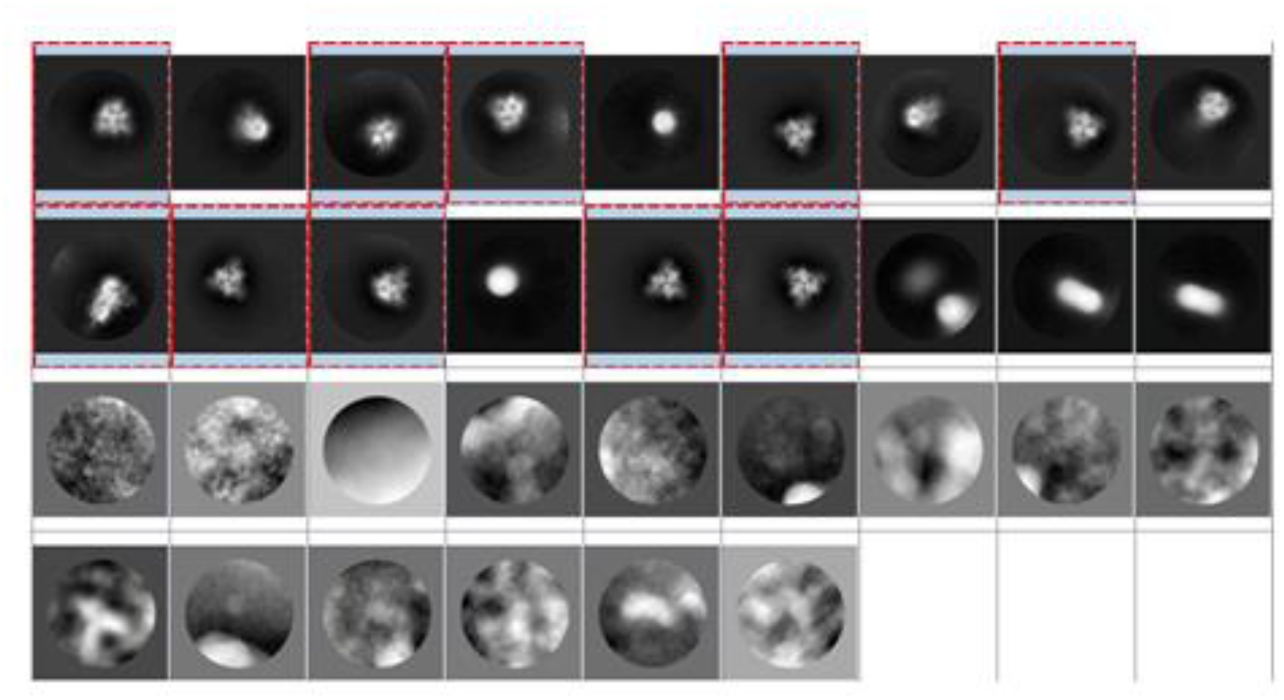
Classification results of SARS-CoV-2 using RELION with 500 set as the prescribed class number.

**Supplementary Fig. 21.**
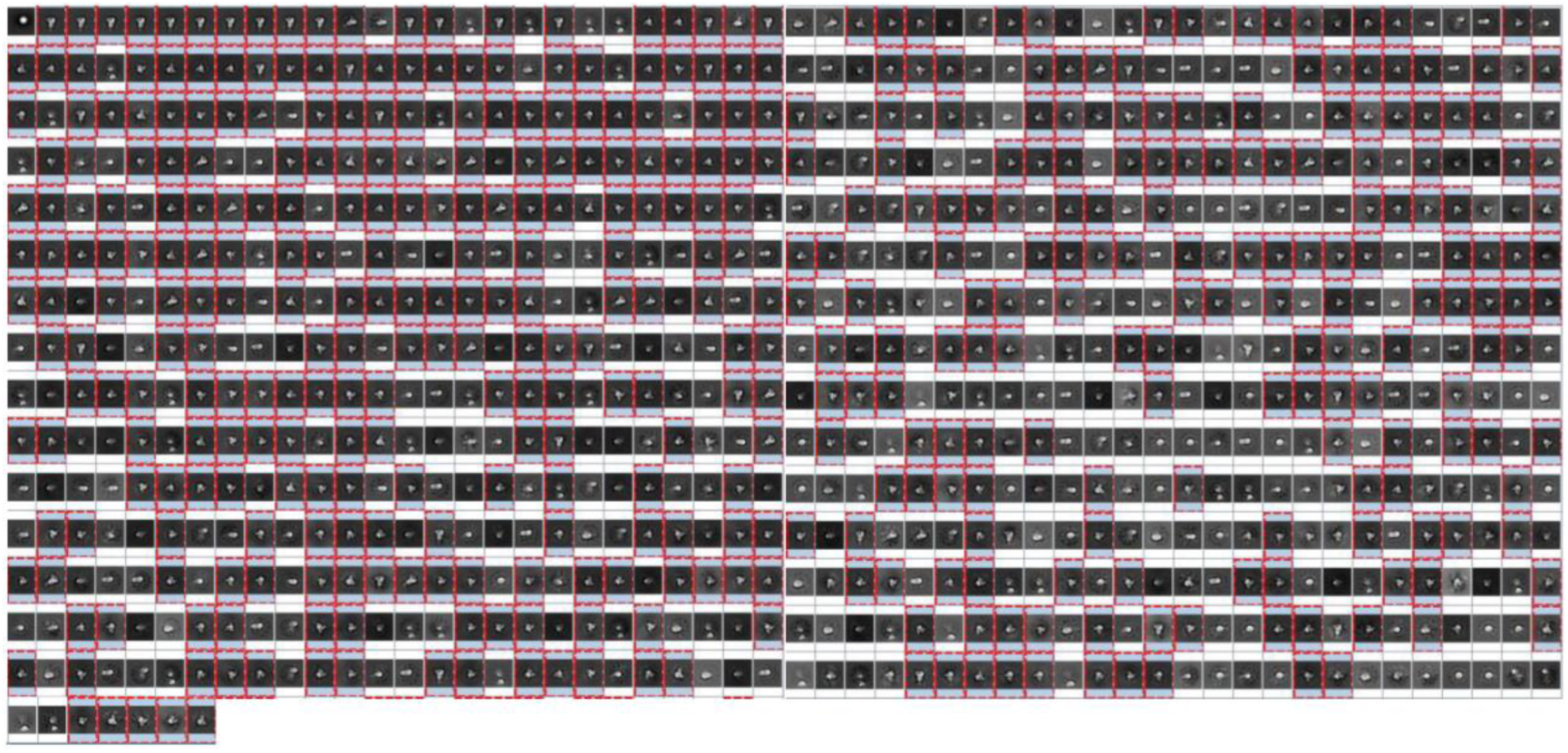
Classification results of SARS-CoV-2 using RE2DC. The high-quality class averages are boxed in red.

**Supplementary Fig. 22.**
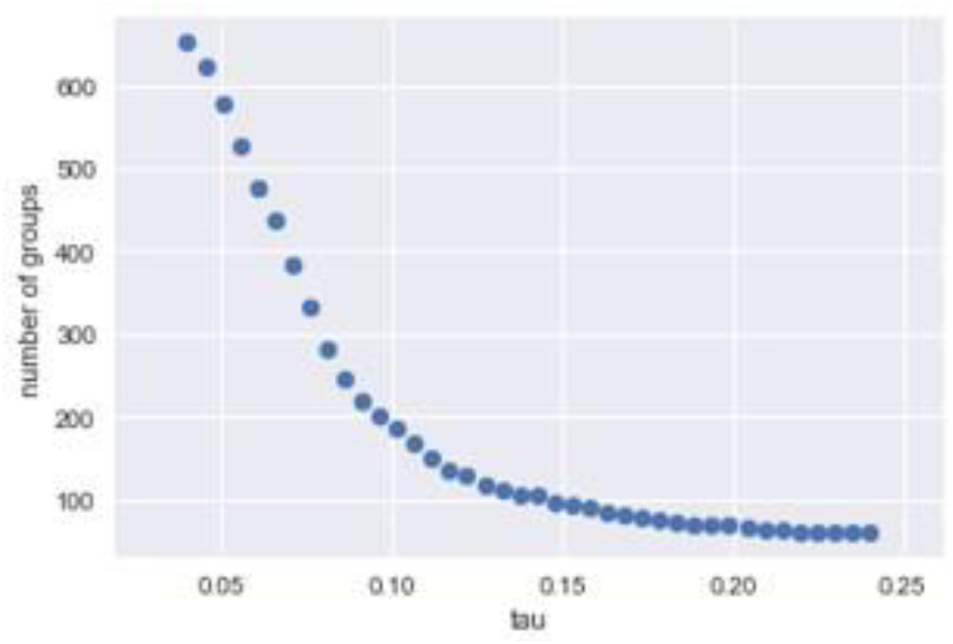
A plot for inspection of the number of clusters generated by γ-SUP under various τ (tau) hyper-parameters for the 80S ribosome dataset. The τ (tau) hyper-parameter for the transition to occur most likely reside in the range of [0.05, 0.075].

## Notes

### Competing Interest Statement

The authors have declared no competing interest.

### Summary of Updates

The main manuscript and Supplementary Information have been extensively revised and reorganized to provide a clearer and more coherent scientific narrative. The associated software implementation has also been updated. The principal results, analyses, and conclusions remain unchanged.

